# FastCCC: A permutation-free framework for scalable, robust, and reference-based cell-cell communication analysis in single cell transcriptomics studies

**DOI:** 10.1101/2025.01.27.635115

**Authors:** Siyu Hou, Wenjing Ma, Xiang Zhou

## Abstract

Detecting cell-cell communications (CCCs) in single-cell transcriptomics studies is fundamental for understanding the function of multicellular organisms. Here, we introduce FastCCC, a permutation-free framework that enables scalable, robust, and reference-based analysis for identifying critical CCCs and uncovering biological insights. FastCCC relies on fast Fourier transformation-based convolution to compute ***p***-values analytically without permutations, introduces a modular algebraic operation framework to capture a broad spectrum of CCC patterns, and can leverage atlas-scale single cell references to enhance CCC analysis on user-collected datasets. To support routine reference-based CCC analysis, we constructed the first human CCC reference panel, encompassing 19 distinct tissue types, over 450 unique cell types, and approximately 16 million cells. We demonstrate the advantages of FastCCC across multiple datasets, most of which exceed the analytical capabilities of existing CCC methods. In real datasets, FastCCC reliably captures biologically meaningful CCCs, even in highly complex tissue environments, including differential interactions between endothelial and immune cells linked to COVID-19 severity, dynamic communications in thymic tissue during T-cell development, as well as distinct interactions in reference-based CCC analysis.

## 1 Introduction

Multicellular organisms rely on cell-cell communications (CCCs) to coordinate development, regulate biological functions, maintain homeostasis, and respond to external stimuli[1–3]. CCCs often occur in the form of ligand-receptor interactions (LRIs), where a ligand released from one cell binds to a receptor on another, triggering downstream signaling events that alter transcription factor activity and gene expression in the receiving cells[3, 4]. These interactions enable cells to coordinate their behavior and facilitate the orchestrated activities of different cell types, driving critical biological processes. Dysregulation of CCCs is frequently associated with pathological conditions, leading to compromised tissue repair, immune dysfunction, cancer progression, and neurodegenerative diseases[2, 5]. Consequently, understanding CCCs is fundamental for unraveling the complex biological processes that drive development and disease.

The widespread availability of single cell transcriptomics data has greatly facilitated the study of CCCs, leading to the development of many computational methods for detecting them[3, 6–15]. These include core tools such as CellPhoneDB(V1[6]), CellChat[9], ICELLNET[10], and SingleCellSignalR[11]; task-specific tools such as CellCall[12] and NicheNet[13] that incorporate intracellular signaling and CellPhoneDB(V4[7] and V5[8]) that expands ligand types; as well as other tools that account for broader conditions, such as CellChat(V1.6 and later), iTALK[14], and Connectome[15], which integrate differential expression analysis to infer CCCs. Despite their differing emphases, these methods all share a common analytic procedure. This procedure begins with a candidate list of LRIs, which can be either user-defined or obtained from existing databases. It proceeds by examining one pair of cell types at a time, calculating a quantity called communication score (*CS*) for each LRI. The value of *CS* is used to determine whether there is coordinated expression of the ligand on cells of one cell type (i.e. senders) and the receptor on cells of the other cell type (i.e. receivers), at levels exceeding what would be expected under the null. Such coordinated expression is suggestive of potential cell-cell communication and interaction.

While the CCC analysis framework is conceptually straightforward, accurate inference of CCCs presents three important challenges. First, determining an appropriate *CS* threshold is technically challenging. Some tools rely on manually setting a default *CS* threshold, which is inherently arbitrary, while others use statistical methods to compute *p*-values, which are essential for rigorously accessing whether the observed coordinated expression exceeds expectations under the null. However, calculating *p*-values is difficult since *CS*, derived from ligand-receptor expression levels, has a relatively complicated functional form that complicates direct evaluation of its null distribution. Consequently, most statistical methods rely on permutation tests to estimate *p*-values. However, permutation approaches are not only time-consuming and computationally demanding but also highly sensitive to the number of permutations performed, affecting usability, accuracy, and reliability of CCC analysis. Second, almost all existing methods rely on a single communication score to evaluate CCC. For example, CellPhoneDB calculates the average expression level of the ligand and receptor, whereas CellChat computes their product. While the average or product can capture specific aspects of CCC, relying on a single score reduces flexibility and limits adaptability to diverse datasets, interaction patterns, and biological contexts. Third, large-scale single cell studies, involving millions of cells, are being conducted, offering unprecedented insights into cellular diversity and function. These studies contain valuable information that could substantially enhance the analysis of CCC. Unfortunately, few methods are capable of scaling to analyze millions of cells efficiently. Moreover, none of the existing methods can take advantage of these large-scale single cell datasets to enhance the analysis of user-collected datasets, which are oftentimes smaller in size, to uncover additional biological insights. These limitations constrain the discovery of critical intercellular signaling pathways and hinder biological understanding of CCCs. To address these challenges, we present FastCCC, a highly scalable, permutation-free statistical toolkit tailored to identify critical CCCs in the form of LRIs and uncover novel biological insights in single-cell transcriptomics studies. FastCCC presents a novel analytic solution for computing *p*-values in CCC analysis, enabling scalable analysis without the need for computationally intensive permutations. It introduces a modular *CS* computation framework that calculates various communication scores through a range of algebraic operations between ligand and receptor expression levels, capturing a broad spectrum of CCC patterns and ensuring robust analysis. Additionally, FastCCC not only enables the analysis of large-scale datasets containing millions of cells, but also introduces, for the first time, reference-based CCC analysis, where large-scale datasets are treated as reference panels to substantially improve CCC analysis on user-collected datasets. To support routine reference-based CCC analysis, we have constructed the first human CCC reference panel, which includes 19 distinct tissue types containing approximately 16 million cells across over 450 cell types, enhancing the interpretability and consistency of CCC analysis across diverse biological contexts. We demonstrate the advantages of FastCCC across multiple datasets, most of which exceed analytical capabilities of existing CCC methods. In real datasets, FastCCC reliably captures biologically meaningful CCCs, even in highly complex tissue environments. Examples include differential interactions between endothelial and immune cells via the C3 ligand and its receptors that are linked to COVID-19 disease severity, dynamic LRIs in thymic tissue during distinct T-cell developmental stages, as well as distinct CCC patterns in reference-based CCC analysis.

## 2 Results

### 2.1 FastCCC Overview

FastCCC is described in the Methods (Section 4), with its technical details provided in the Supplementary information and its core method schematic shown in Fig 1. FastCCC provides three major advances. First, FastCCC presents a novel, alternative strategy for computing *p*-values in CCC analysis. By leveraging convolution techniques through Fast Fourier Transform (FFT), FastCCC derives the analytic solution for the null distribution of *CS*, enabling scalable analytic computation of *p*-values without requiring computationally intensive permutations. This approach makes FastCCC orders of magnitude faster than existing methods, while enhancing accuracy and robustness. The computational gain of FastCCC further increases with larger datasets, making it particularly suited for large-scale, state-of-the-art single cell studies that existing methods cannot handle. Second, FastCCC introduces a modular *CS* computation framework that captures interaction strength through algebraic operations between ligand and receptor expression level. This framework enables the development of new *CS* scores beyond common ones by utilizing a wide range of expression summary statistics to capture a broad spectrum of CCC patterns. It also accommodates multi-subunit protein complexes for ligands and receptors, accounting for distinct interaction strength characterized by different subunits. As such, FastCCC substantially enhances the power and robustness of CCC detection across diverse datasets and biological contexts. Finally, FastCCC not only offers a scalable and effective solution for CCC analysis, enabling the analysis of large-scale datasets containing millions of cells, but also, for the first time, allows these large-scale datasets to serve as reference panels, facilitating more comprehensive and context-aware CCC analysis on user collected datasets, which may be much smaller in size. To support routine reference-based CCC analysis with FastCCC, we constructed the first human CCC reference panel with 19 distinct tissue types and approximately sixteen million cells (Methods, Table A1). This reference panel enhances the interpretability and consistency of CCC analysis across diverse biological contexts.

**Fig. 1.**
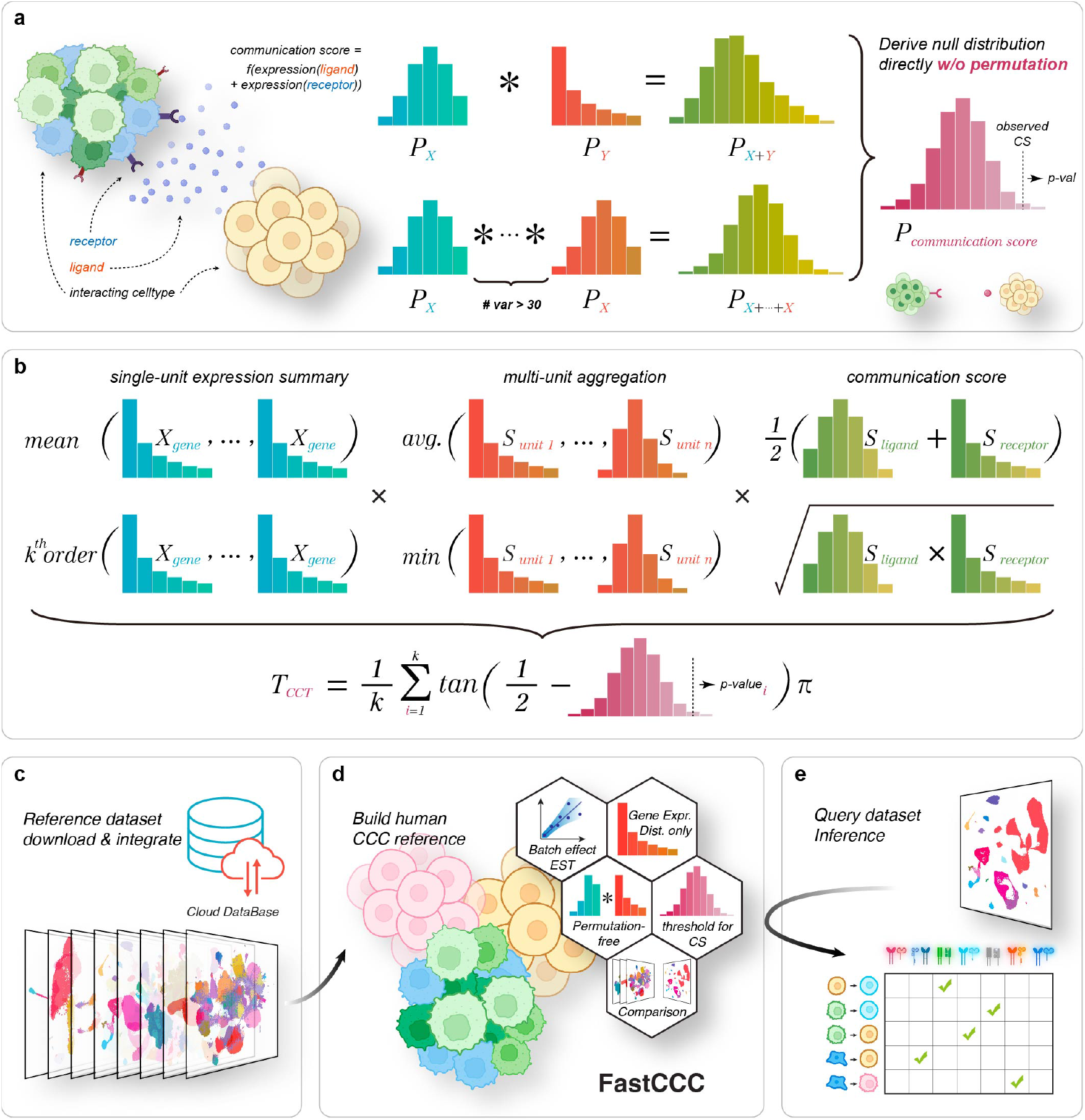
Overview of FastCCC algorithm. **a**. A simplified diagram illustrating the core idea of the FastCCC workflow. Fast Fourier transform-based convolutions are employed to derive an analytic solution for the null distribution of various cell-cell communication scores, which are calculated through extensive summation, averaging, and logarithmic summation of multiplication operations. **b**. The modular scoring system in FastCCC enables the computation of diverse communication scores (*CS*), utilizing either mean or *k*-th order statistics for single-unit expression summaries and various aggregation methods such as average or minimum. Users can freely combine different modules for *CS* calculation. When the option of multiple *CS* combinations are selected, FastCCC also provides a Cauchy combination test to integrate *p*-values from various modular scores. **c**. Single-cell RNA-seq transcriptomic data from a wide range of large-scale human normal tissue samples were processed and integrated to create the first human CCC reference panel. **d**. FastCCC was used to create the reference panel by calculating gene expression distributions and key statistical metrics on large-scale single-cell sequencing data. **e**. The constructed reference panel is used for CCC inference in the query data. In particular, FastCCC performs reference-based CCC analysis in the user-provided query dataset by leveraging the constructed human CCC reference panel to identify key CCCs in the query data.

### 2.2 FastCCC ensures accurate and scalable *p*-value computation

The first important feature of FastCCC is its ability to compute *p*-values analytically, rather than relying on computationally intensive permutations. To validate the analytic solutions provided by FastCCC, we first focus on the same *CS* statistic used in CellphoneDB (CPDB), and compared the *p*-values from FastCCC, calculated using this single *CS* statistic, with those from CPDB, which uses permutations. We began by testing the tutorial data used in CPDB (referred to as CPDBTD), which comprises 3,312 cells across 40 cell types, nearly half of which are rare, with fewer than 30 cells per cell type, including five extremely rare cell types with fewer than ten cells (Fig 2a). The cell type composition in the data captures a range of the strategies employed by FastCCC to compute *p*-values (see Methods; Table A2). As expected, the *p*-values from FastCCC are highly concordant with those obtained from CPDB (Pearson correlation *>* 0.995), with the consistency increasing as the number of permutations increases (Fig A1). For example, when evaluating the set of significant LRIs (*p*-value *<* 0.05), the intersection over union (IoU) between FastCCC and CPDB, using the default CPDB setting of 1,000 permutations, is 96.7%, with precision exceeding 99.3%. These metrics continue to improve as the number of CPDB permutations increases (Fig 2b, A2, A3): with one million permutations, Pearson correlation increased to 0.988 (Fig 2c), while IoU and precision reach 97.3% and 99.7%, respectively. Additionally, the correlation for valid LRIs, defined as those where both the non-zero expressed ligand in the sender and the receptor in the receiver exceed a specified percentile, was 0.996 (Fig A4, Methods). Treating the results of CPDB with one million permutations as ground truth, we found that using 1,000 to 50,000 permutations in CPDB resulted in a notable drop in precision and IoU metrics, particularly under stringent *p*-value thresholds (Fig A5). In contrast, the analytical solution provided by FastCCC maintained high accuracy. Notably, CPDB requires at least 10,000 permutations – ten times its default setting – to match FastCCC’s *p*-value accuracy. Importantly, we also modified CPDB’s *CS* formulate and validated FastCCC’s *p*-value results across a range of *CS* using operations, including order statistics and geometric mean (Fig A6, Table A2, and Methods).

**Fig. 2.**
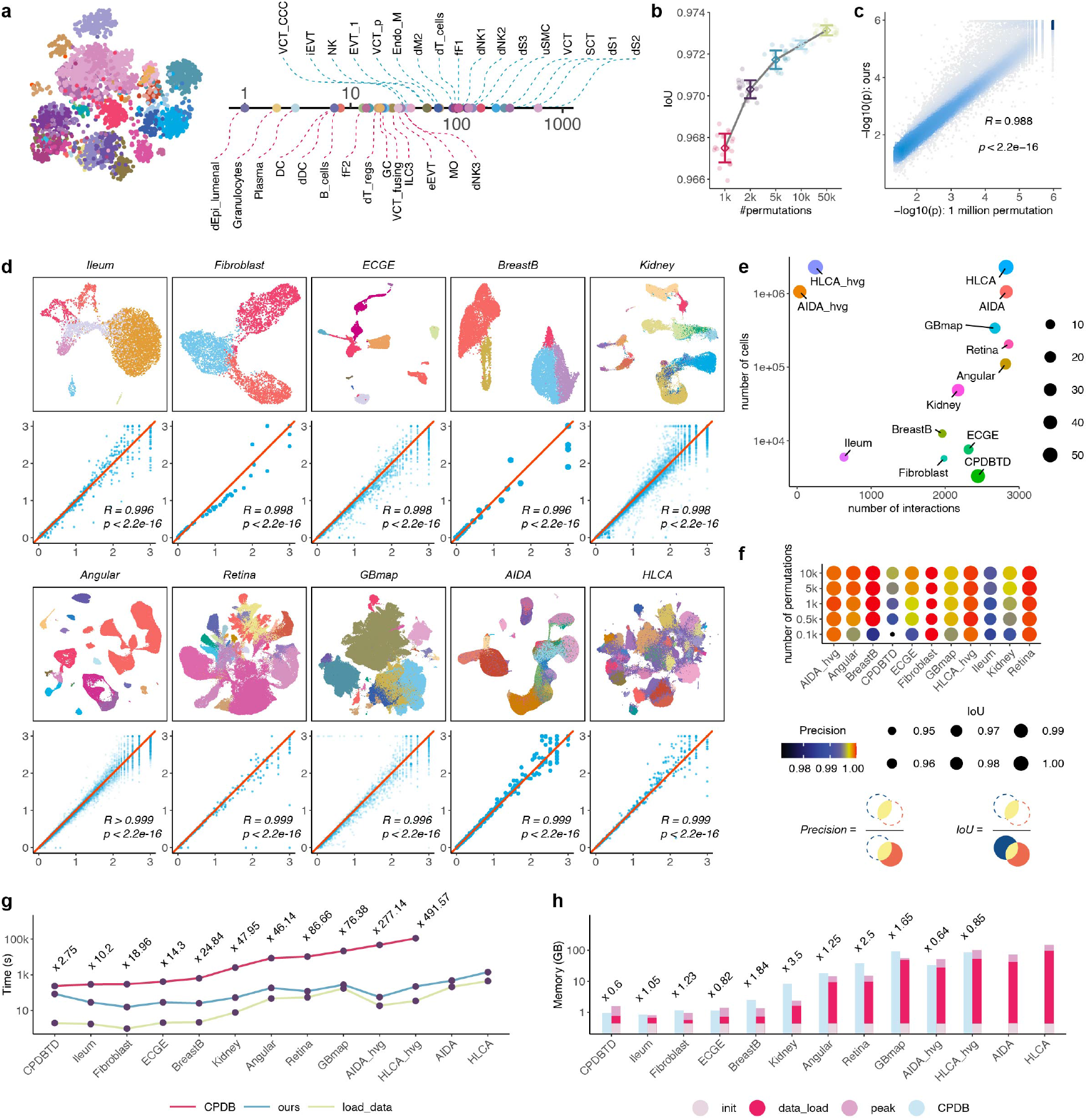
Analytical assessment of FastCCC. **a**. UMAP plot of CPDBTD (left) and a line plot shows the corresponding cell counts for each cell type (right). The pink dashed line indicates rare cell types with fewer than 30 cells while the blue dashed line indicates common cell types. **b**. IoU results on the significant LRIs detected between FastCCC and CPDB across varying numbers of permutation tests used in CPDB. Each circle represents an individual experiment (20 replications), with squares and error bars indicating the mean and variance. **c**. Comparison of FastCCC results with CPDB (1 million permutation tests). Each point represents a significant LRI identified by CPDB. The x-axis shows CPDB p-values on -log10 scale, and the y-axis shows FastCCC results. Pearson correlation coefficient and corresponding p-value are annotated. **d**. UMAP plot of ten additional scRNA-seq datasets, with different colors representing different cell types. The p-value for valid LRIs between FastCCC and CPDB is compared, with Pearson correlation coefficient and corresponding p-value annotated. **e**. Scatter plot displays the number of cells and the number of interaction types in each dataset. **f**. Precision and IoU results for different numbers of permutation tests across all test datasets. **g**. Running time of CPDB and FastCCC, along with data loading time, across all test datasets. **h**.Memory consumption of CPDB and FastCCC, along with the memory used for data loading, across all test datasets.

Next, we assessed the consistency of FastCCC with CPDB across ten additional scRNA-seq datasets, encompassing a wide range of cell type numbers, dataset sizes, and ligand-receptor pairs (Fig 2e, Methods). A high level of consistency was again observed between the *p*-values from the two methods, with Pearson’s correlation ranging from 0.996-0.999, confirming the reliability of FastCCC (Fig 2d). Moreover, the consistency between the two methods improves with an increasing number of permutations used in CPDB, with precision surpassing 99% and IoU surpassing 97% across all datasets when the number of permutation reached 10,000 (Fig 2f).

While the *p*-values from FastCCC are nearly identical to those from CPDB, FastCCC offers significant computational advantages in terms of both computation time and memory usage. These efficiency gains brought by FastCCC increase as the number of cells in the dataset grows. Specifically, FastCCC is at least 40 times faster than CPDB when the number of cells exceeds 100,000 and nearly 500 times faster on a dataset with two million cells, completing all calculations under 20 minutes in the latter case (Fig 2g). In fact, the computation of FastCCC is so efficient that a substantial proportion of its required computing resources, approximately 60% of memory and 25% of computing time across all test datasets, are spent on data reading rather than the actual computation (Fig 2g,h). Additionally, across the eleven smaller datasets, the average memory consumption of FastCCC was 69% of CPDB; while FastCCC is the only method that is scalable to the two large datasets (AIDA and HLCA), each containing over one million cells with thousands of different LRIs.

We extended the computation comparison to the default version of FastCCC, which uses 16 distinct *CS* values (more below), along with seven existing CCC analysis methods, including CellphoneDB V5 and V4, CellChat, SingleCellSignalR, ICELLNET, NicheNet, and CellCall. These methods were selected based on their superior performance in previous benchmarking studies[16, 17] and their widespread usage in the field. Compared to these approaches, FastCCC stands out as the only method that provides statistical significance without the need for permutations (Fig 3a). For the eight smaller datasets, FastCCC was at least 10 times faster than nearly all other methods, while also consuming the least memory across datasets (Fig 3b,c). For the two large datasets, FastCCC was again the only applicable method. In particular, CellChat and CellCall encountered memory errors when processing datasets with 100k-200k cells using default parameters, while SingleCellSignalR timed out when processing datasets over 100k cells (see Methods). Both NicheNet and CellCall exhibited significant delays and timeouts on datasets larger than 500k cells. CellPhoneDB series showed excessive memory consumption on datasets reaching 1 million cells. In contrast, FastCCC efficiently processed large-scale datasets with high speed and low memory consumption, demonstrating its scalability and practical application in real-world scenarios involving massive scRNA-seq datasets.

**Fig. 3.**
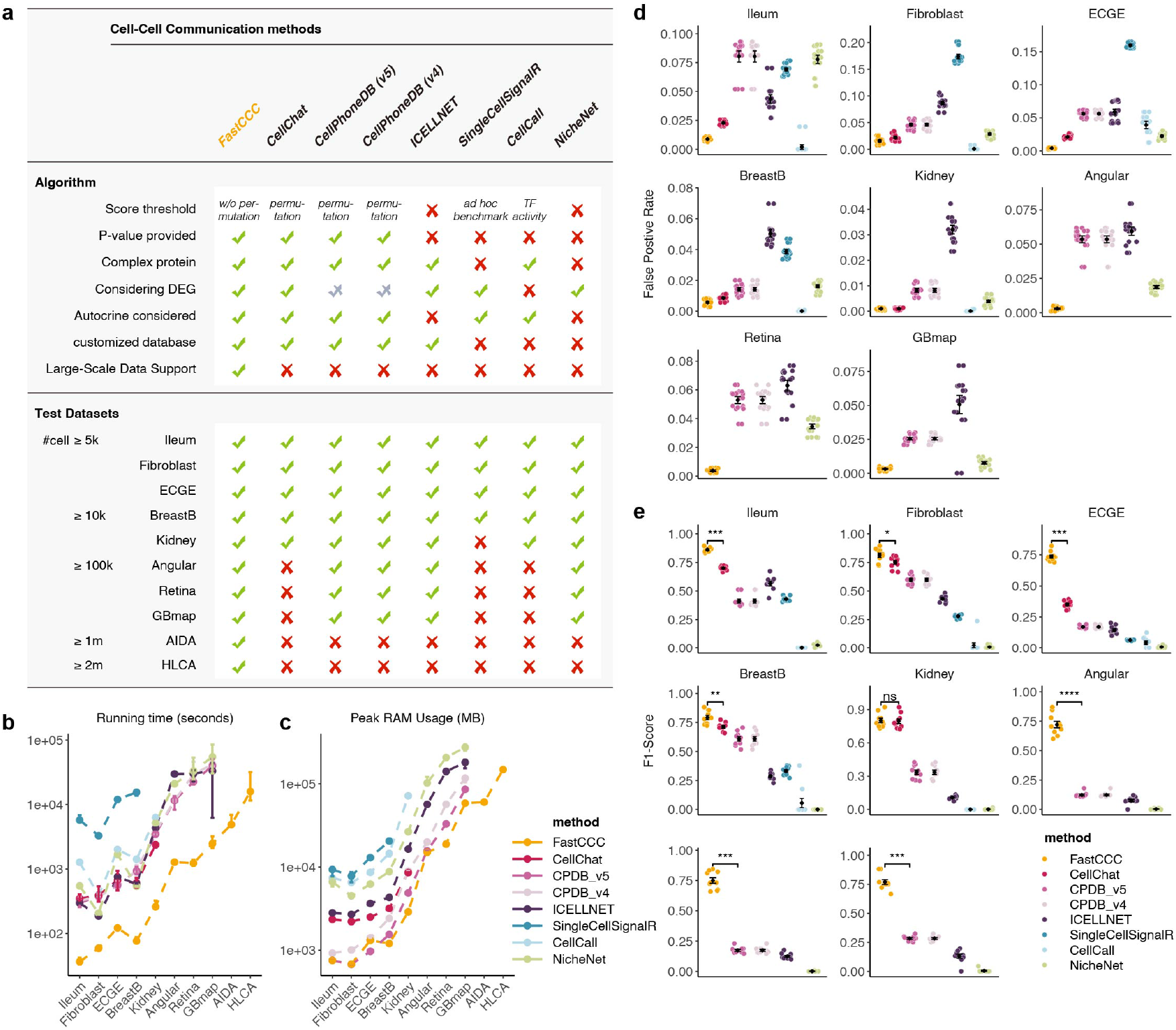
Performance comparison of FastCCC with other methods. The results are shown for the default version of FastCCC with 16 scoring methods **a**. A comparative table outlining the key features, functionalities, and applicability of different CCC analysis methods across test datasets. ✓indicates the presence of a feature or compatibility. *×* indicates the absence of a feature or incompatibility. Half tick for CPDB in some features because it requires users to provide DEG data. **b**. FastCCC vs. other methods in runtime across 10 datasets. Datasets are sorted by the number of cells on the x-axis. Runtime is shown on the y-axis in seconds. **c**. FastCCC vs. other methods in memory consumption across 10 datasets. Peak memory usage is shown on the y-axis in MB.**d**. FastCCC vs. other methods in terms of FPR across 8 test datasets. Each point represents an individual simulation experiment. The mean and standard error are annotated. Lower values (closer to 0) indicate better performance. **e**. FastCCC vs. other methods in terms of F1-score across 8 test datasets. Each point represents an individual simulation experiment. The mean and standard error are annotated. Higher values (closer to 1) indicate better performance.

### 2.3 FastCCC is robust and powerful

The second important feature of FastCCC is its ability to employ a variety of algebraic operations to compute additional *CS* values beyond those previously used, such as the one used in CPDB. The default version of FastCCC includes a total of 16 *CS* values, allowing it to capture a wide range of interaction patterns between ligands and receptors across different biological contexts, thus ensuring its power and robustness. Here, we evaluated the accuracy of FastCCC and compared it with existing methods on eight relatively small single cell datasets (excluding AIDA and HLCA), where most other methods were able to produce results. To achieve this, we employed a semi-simulated approach: shuffling cell labels to create null scenarios where none of the LRIs interact with each other, and a small proportion of LRIs were then randomly selected and overexpressed to serve as the truly interacting LRIs, thereby establishing ground truth (see Methods).

We first examined false positive rates (FPR) and recall of various methods across datasets. Nearly all methods achieved FPR below 10% across all datasets (Fig 3d), with the only exception of SingleCellSignalR, which exhibited an average of 17.4% and 16.0% FPR in the Fibroblast and ECGE datasets, respectively. FastCCC achieved the lowest FPR among all methods, maintaining FPR below 1% across all datasets, except Fibroblast, where the FPR was 1.5%. CellCall also achieved a relatively low FPR, but its recall was nearly zero across all datasets (Fig A7), suggesting that it failed to detect any signals. Similarly, NicheNet also yielded low recall. In contrast, FastCCC achieved the highest recall while maintaining the lowest FPR, reflecting its superior accuracy in capturing the true interactions.

Beyond recall and FPR, FastCCC consistently outperformed the other methods across all datasets on other evaluation metrics such as precision, accuracy, specificity, balanced accuracy, and the Jaccard index (Fig A7). Additional metrics, such as F1 score and Matthews correlation coefficient (MCC), also yielded similar conclusions (Fig 3e, Fig A7), supporting FastCCC’s superior performance. For instance, in the Ileum dataset, FastCCC achieved the highest median and mean F1 scores, both at 0.863, outperforming CellChat (median: 0.699, mean: 0.701), CellPhoneDB V5/V4 (median: 0.391, mean: 0.413), SingleCellSignalR (median: 0.425, mean: 0.431), ICELLNET (median: 0.574, mean: 0.565), NicheNet (median: 0.027, mean: 0.025), and CellCall (both 0.0 due to no significant output). Notably, among the other methods, methods employing *p*-values, such as CellChat and CellPhoneDB, generally outperformed others, which aligns with previous benchmarking studies[16, 17], highlighting the value of rigorous statistical approaches in CCC analysis.

### 2.4 Application of FastCCC to a large-scale COVID-19 dataset

The scalability and effectiveness of FastCCC allows us to conduct CCC analysis on large-scale single-cell datasets within a reasonable time frame and at a fraction of computational cost of the other methods. To further illustrate its utility, we applied FastCCC to two large-scale datasets: a large COVID-19[18] dataset, and a thymus dataset[19]. The second data contains pseudo-time information that requires CCC analysis to be conducted across time points.

We first applied FastCCC to a COVID-19 scRNA-seq dataset comprising 1.46 million cells[18] (Fig 4a), a scale beyond the feasible resource limits of other comparison methods. FastCCC successfully completed all 16 distinct score calculation methods in approximately 40 minutes, using 120 GB of memory (with the dataset itself occupying 60 GB). This dataset includes samples from 40 patients and 5 healthy controls, categorized into different stages – disease progression and recovered convalescence (Fig 4b). The dataset also contains epithelial (Epi) cells and various immune cells sampled from patients with moderate and severe disease. We explored significant CCCs across different groups to understand changes in cellular communication during COVID-19 progression.

**Fig. 4.**
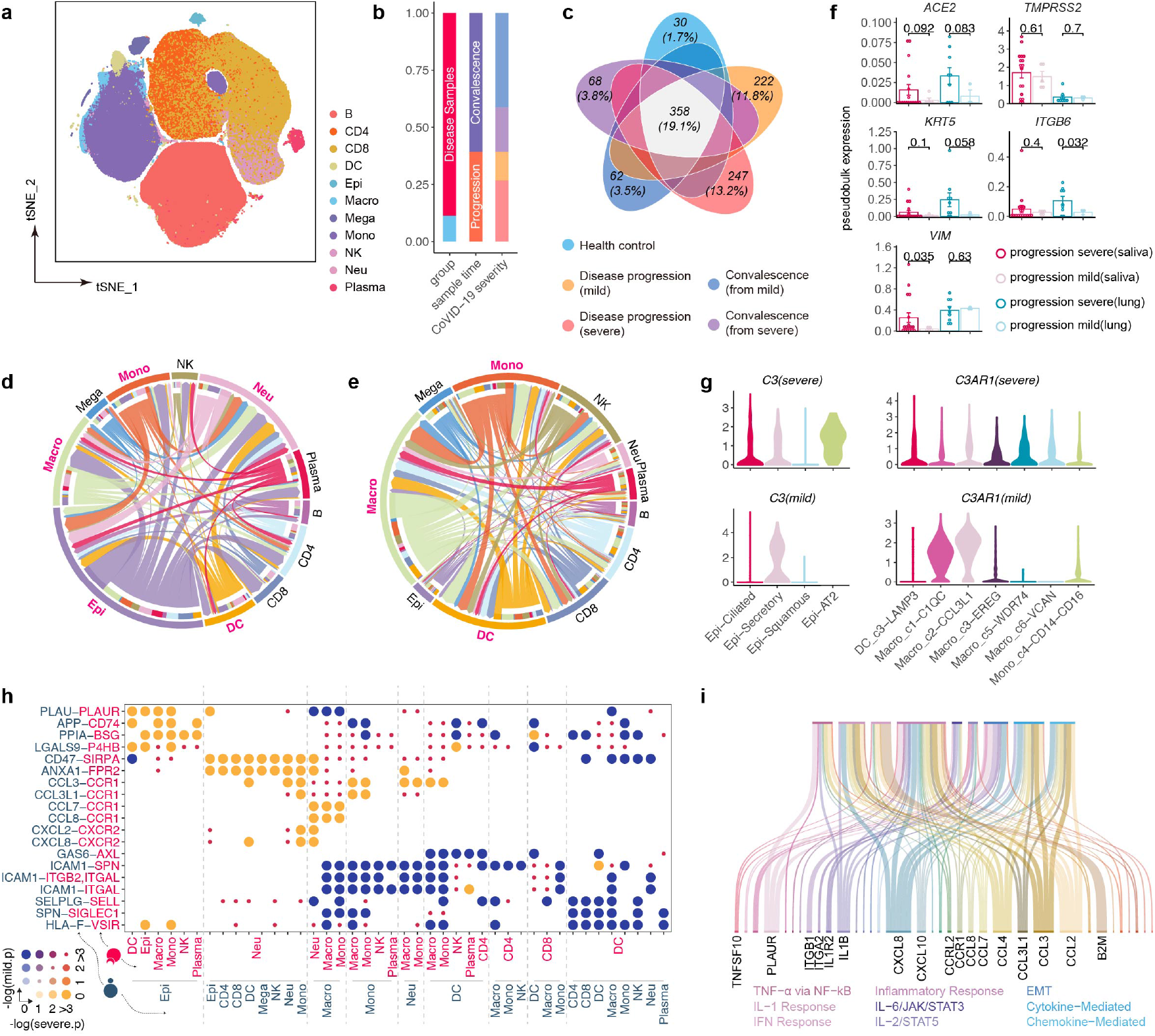
Application of FastCCC to a large COVID-19 dataset. **a**. UMAP plot of the COVID-19 dataset, with colors representing different cell types. **b**. A percent stacked bar chart showing sample groups, disease sampling time points, and COVID-19 severity levels. **c**. Venn diagram of significant LRIs across different groups. The number and proportion of unique LRIs in each group are annotated, with additional labels for intersections comprising more than 10%. **d**. Chord plot of significant LRIs uniquely expressed in the progression severe group compared to the mild group. The color and angle of the arcs represent cell types and the number of significant LRIs involved, respectively. The color of the chords indicates the sender cell type, with the width reflecting the number of LRIs. Arrows point to the receiver cell type, and the main cell types involved in CCC are annotated. **e**. LRIs uniquely expressed in the mild group compared to the severe group. **f**. Pseudo-bulk RNA-seq expression of signaling molecules involved in CCCs during disease progression compared to healthy controls. Each point represents a sample, with colors indicating different sample sources and severity. Mean and standard deviation for each group are indicated, and *p*-values between groups are annotated. **g**. Violin plot showing the expression levels of ligand C3 and its receptor C3AR1 across different groups. **h**. Circle plot of differentially expressed LRIs. The size and color of the circles together represent the significance of LRIs in the mild and severe groups. **i**. Pathways associated with the ligands and receptors differentially expressed in the severe group compared to the mild group are displayed.

We first conducted CCC analysis across five sample groups, including healthy controls, patients with mild and severe cases during disease progression, and patients in recovery, categorized as mild and severe convalescence. FastCCC detected a total of 1,874 CCCs by integrates 16 *CS* scores, with 702, 1,273, 1,026, 996, and 992 CCCs identified in the five groups, respectively (Fig A9). A significant proportion of CCCs (358 CCCs, 19.1%) were shared across all groups, with 7-170 intersections (mean=45.4; median=24) shared between pairs of groups (Fig A9). Importantly, we observed a progressive divergence from the healthy control group, moving through mild recovery, severe recovery, mild disease, to severe disease (Fig A10), as reflected by the increasing number of unique CCCs detected in these groups. For instance, the healthy control group exhibited only 30 unique CCCs, while the convalescence groups showed approximately 60 unique CCCs. In contrast, the disease progression groups displayed over 200 unique CCCs (Fig 4c). Additionally, 9.1% (170) of CCCs were shared exclusively between the mild and severe disease progression groups.

We narrow down our focus to examine the CCCs identified by FastCCC in severe and mild COVID-19 case to interrogate the possible molecular pathways underlying disease progression. We found that the CCCs uniquely occurring in severe cases primarily involved interactions among epithelial cells and myeloid cells, such as macrophages (Macro), neutrophils (Neu), monocytes (Mono), and dendritic cells (DCs)(Fig 4d), In contrast, CCCs occurred in mild cases primarily involved interactions between Macro, DCs, Mono (Fig 4e), suggesting a different pattern of immune response involvement during COVID-19 progression (Fig A10). The top signaling molecules involved in CCCs with disease progression compared with health control include ACE2, TMPRSS2, KRT5, ITGB6, and VIM emerged as significant ligands or receptors in patients. ACE2 and TMPRSS2 are key components of the COVID-19 pathway[20–22], with ACE2 serving as the functional receptor for the spike glycoprotein of SARS-CoV-2, while TMPRSS2 serving to cleave the spike protein, facilitating viral entry into host cells. To further investigate, we obtained pseudobulk RNA expression data for each sample and focused on epithelial cells from two primary sources: saliva and lung, capturing nasal epithelial cells and upper airway epithelial cells, respectively. We examined the differential CCC information to explore possible communication mechanisms involving these epithelial cells. We found that (Fig 4f), while both ACE2 and TMPRSS2 are significantly elevated in COVID-19 patients compared to healthy controls, their expression patterns differ between severe and mild cases: ACE2 is marginally more expressed in severe cases, both in nasal and upper airway epithelial cells, whereas TMPRSS2 expression does not differ significantly between the two groups. Additionally, markers of epithelial-mesenchymal transition (EMT), such as KRT5, ITGB6, and VIM, were associated with COVID-19 severity. Specifically, KRT5 and ITGB6 were more highly expressed in lung tissues of severe patients compared to mild cases, while VIM was highly expressed in the upper airway of severe patients but showed no significant difference in lung tissues. Therefore, epithelial cells in severe cases appear to secrete ligands that enrich EMT pathways compared to mild cases, dovetailing recent studies and suggesting significant differences in disease progression mechanisms between severe and mild cases[23, 24].

Another top signal identified during CCC analysis is the ligand C3, which directs the interaction between Epi cells and various immune cells such as Macro, Mono, and DCs. While C3 is typically produced in the liver and also secreted by adipocytes and macrophages as a central component of the immune system[25], it displayed an unusually highly expression in ciliated, secretory, and AT2+ epithelial cells in the severe group(Fig 4g). In mild cases, C3 was only highly expressed in secretory epithelial cells, which constitute less than 1% of the total epithelial cells. The results suggest that C3 activation drives maladaptive immune responses, promoting the release of pro-inflammatory cytokines and recruitment of neutrophils and monocytes into lung tissue[26–29]. These immune cells can damage the air–blood barrier, thereby exacerbating ARDS severity in SARS-CoV infections[26].

Other signals differentially detected between severe and mild cases include chemotaxis, such as when Epi cells release PPIA as a ligand to bind with BSG (CD147) receptor, directing Macro, Mono, NK cells and others to the site of infection or inflammation. Additionally, severe cases showed elevated expression of CCCs related to chemokines like CCL and CXCL (Fig 4h). These findings imply the occurrence of a cytokine storm in severe patients, contributing to their worsened clinical phenotypes. In contrast, mild patients exhibited higher expression of leukocyte adhesion-related CCCs, primarily between innate immune cells.

Finally, pathway enrichment analysis of genes involved in these differential CCCs revealed that severe patients activate multiple immune pathways (e.g., TNF*α* via NF-*κ*B, IL-6/JAK/STAT3), which are likely contribute to cytokine and inflammatory storm formation(Fig 4i, Fig A11, A12).

Applying our method to this large-scale dataset demonstrated its practical utility, with the COVID-19 results largely corroborating existing research while uncovering numerous unique signals that may offer new avenues for further study.

### 2.5 Application of FastCCC to a thymus dataset with developmental time series

We applied FastCCC to a large-scale thymus scRNA-seq dataset with pseudo-time information, dividing it into thousands of sliding time windows for detailed characterization. This required CCC anlaysis to be carried out within each time window, resulting in thousands of analyses—a burdensome task that again is not feasible by other methods—to comprehensively investigate the dynamic cellular interactions during T cell development. This data consisted a total of 255,901 cells[19], including thymic epithelial cells (TECs), dendritic cells (DCs), various stromal cells, as well as differentiating T cells (Fig 5a). The T cell population consists of 76,903 thymocytes at different developmental stages, including double negative (DN), double positive (DP), CD4+ single positive (SP), CD8+ SP, and T regulatory (Treg) cells, among others.

**Fig. 5.**
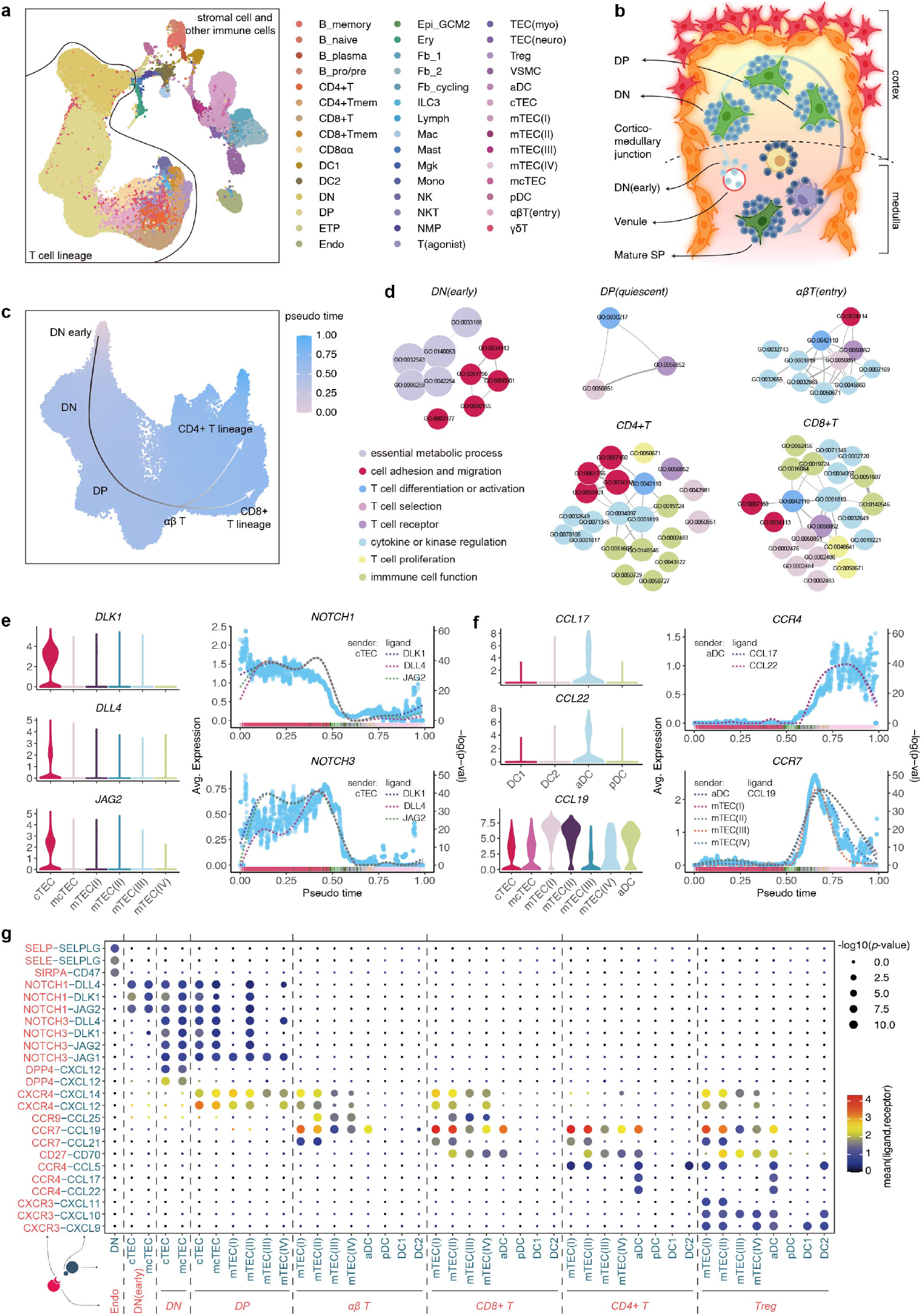
Application of FastCCC to the developmental thymus dataset. **a**. UMAP plot of the thymus dataset, with colors representing different cell types. Developmental T cell lineage and other stromal cells or immune cells are separated by dividing lines. **b**. Illustration of the T lymphocyte developmental process. **c**. UMAP plot of the T cell lineage pseudotime, showing progression from the earliest DN stage to CD4+ or CD8+ lineages. **d** GO pathway enrichment analysis of signaling molecules involved in key CCCs at each stage. **e**. Violin plot showing expression levels of ligand DLK1, DLL4, and JAG2 across different stromal cell type groups. The corresponding receptor expression levels (NOTCH1/3) and *CS p*-value over time are depicted, where each blue dot represents the mean receptor expression within a time window, and dashed lines indicate the smoothed -log(p-value) levels for a specific LRI pair. **f**. Violin and scatter plots showing expression levels of ligands CCL17/19/22, along with their corresponding receptors CCR4/7, as well as the changes in *p*-value. **g**. Circle plot of significant LRIs between stromal cell types and thymocytes at different developmental stages.

Thymic development is a highly regulated process in which T lymphocytes originate from multipotent hematopoietic stem cells located in the bone marrow[30]. Initially, progenitor cells migrate from the bone marrow to the thymus via the blood, entering through the cortico-medullary junction before moving to the outer cortex. Within the thymus, these progenitors undergo a series of maturation steps[19, 30]. As shown in the Fig 5b, the thymus is composed of multiple lobules, each containing a distinct outer cortical region and an inner medulla. Developing T-cell precursors reside within a complex epithelial network known as the thymic stroma, a specialized microenvironment crucial for T-cell development. This stroma is rich in signaling pathways that regulate the commitment to the T-cell lineage.

Upon entering the thymus, progenitor cells lack the surface molecules characteristic of mature T cells, and their receptor genes remain unrearranged. These progenitor cells, often referred to as “double-negative” thymocytes due to their lack of CD4 and CD8 expression, eventually differentiate into the major population of *αβ* T cells and the minor population of *γδ* T cells. Further differentiation of *αβ* T cells results in the formation of two distinct functional subsets: CD4+ T cells and CD8+ T cells. Our study primarily focuses on the major *αβ* T-cell lineage. By analyzing thymocytes as a whole, we examined the interactions between these cells with other stromal cells and immune cells within the thymic microenvironment.

We first examined the relationship between pseudotime and T cells types (Fig 5c) and found that the inferred pseudotime trajectory captures the expected T cell development from early thymic progenitors (DN (early)), to DP cells, and eventually to either CD8+ or CD4+ SP T cells, which further specialized into subsets including T helper cells (Th) and regulatory T cells (Tregs). Next, we conducted differential expression analysis using FastCCC’s probability toolkit for each T cell subpopulation and performed pathway enrichment analysis on significant ligands and receptors identified at various developmental stages (Fig 5d). These confirmed ligands and receptors were involved in CCCs that revealed the critical molecular pathways underlying T cell development. Specifically, during the DN stage, T-cell communication predominantly involved essential metabolic processes related to cell survival, proliferation, and development. However, in the DP (quiescent) stage, the number of significant interactions decreased sharply, with the remaining interactions strongly associated with T-cell differentiation, activation, and selection, indicating the onset of T-cell lineage commitment. In the *αβ* T-cell stage, additional cytokine and kinase regulation pathways emerged, along with signals related to cell adhesion and migration, suggesting that T cells were beginning to mature and migrate out of the cortex. Once T cells differentiated into CD4+ and CD8+ subsets, the signaling landscape became more complex, with an increase in interactions related to immune cell functions and T-cell proliferation. Signals associated with cell adhesion and migration also reappeared, indicating that these mature T cells were preparing to exit the thymus and circulate in the periphery.

FastCCC identified multiple key signals during T cell development, including NOTCH and chemokine signaling pathways. First, FastCCC accurately depicted the intense interactions driven by NOTCH1 and NOTCH3 receptors and their ligands, such as DLK1, DLL4, and JAG2, during the DN(early) and DN stage (Fig 5e). NOTCH receptor signaling is key for T-cell precursors commitment to the T-cell lineage[30]. Second, FastCCC identified multiple chemokine receptors such as CCR4 and CCR7, along with their ligands (e.g., CCL17, CCL22, CCL19), to serve as key players during the migration of post-selection thymocytes into the medulla (Fig 5f). This result implies the role of chemokine signaling in guiding thymocyte migration to the medulla and ensuring the proper maturation of SP thymocytes. In particular, we observed that CCL19 and CCL21 released from mTECs (medullary TECs) interacted with CCR7 receptor on naïve lymphocytes to potential controlling the migration of developing thymocytes. Such interaction between CCR7 and its ligands started in the *αβ* T-cell (entry) stage and the interaction strength peaked during the SP stage.

Among the CCR7 ligands, CCL19 is specifically expressed in mTECs compared to cTECs (cortical TECs), and such expression pattern between mTECs and cTECs likely created a chemotactic gradient that guides the migration of CCR7-bearing naïve T cells[31–34]. Interestingly, we found that a subtype of dendritic cells (aDCs) also expressed ligands such as CCL17, CCL19, and CCL22 (Fig 5f,g), suggesting that this specific DC subtype, but not other DC subtypes such as pDCs, DC1, and DC2, may play a key role in modulating the thymic niche.

FastCCC identified multiple additional CCCs that also play crucial roles during T cell development (Fig 5g). For instance, FastCCC revealed that P-Selectin Glycoprotein Ligand (SELPLG or PSGL-1), a glycoprotein characterized by mucin-type O-glycan modifications that are expressed in DN cells, interacts with E- and P-selectins (SELP, SELE) on TECs. This interaction mediates cell rolling and tethering, promoting the capture of cells from the bloodstream, providing support that endothelial cells assist T-cell progenitors in entering the thymus during the colonization of lymphoid progenitor cells[31]. Additionally, FastCCC also detected CD47-SIRPA signaling, which promotes the opening of tight connections between endothelial cells, to mediate the communication between DN and endothelial cells[31, 35]. Regarding CCL21 and CCR7 signaling, similar to the CCL19-CCR7 signal, we discovered that CCL21 is expressed in both type I and type II mTECs, while CCL19 is expressed across all mTEC subtypes. Our method also identified that the CXCR4 receptor and its ligand CXCL12, as well as the CCR9 receptor and its ligand CCL25, are vital signals directing thymocyte migration.

These findings illustrate that thymocytes are not merely passive participants within the thymus; they actively influence the organization of the thymic epithelial cells on which they depend for survival. As developing thymocytes progress through distinct stages, marked by changes in T-cell receptor expression, these surface changes reflect their functional maturation. Specific combinations of cell surface proteins serve as markers for T cells at different stages of differentiation. Collectively, our results suggest that FastCCC effectively captures significant signaling interactions across various developmental stages, providing invaluable insights into the temporal dynamics of these interactions.

### 2.6 Application of FastCCC for reference-based CCC analysis and construction of a human CCC reference panel

The third important feature of FastCCC is its ability to conduct reference-based CCC analysis. With the growing availability of large-scale single-cell RNA sequencing datasets across diverse tissues and samples[36], FastCCC can utilize these datasets as reference panels. By leveraging the information contained in the reference panels, FastCCC enhances CCC analysis on user-collected data, which are often smaller in size, yielding more comprehensive and insightful results.

To support reference-based CCC analysis, we first constructed a human CCC reference panel using the CELLxGENE[37] platform. Specifically, we curated and organized 16 million singe cell data from 19 human tissues, containing 452 cell types from 1,025 healthy samples and 10 distinct techniques(Fig 6a,b, and Table A1). This effort resulted in the creation of the first human CCC reference panel, which can be used by FastCCC to directly identify significant CCCs by comparing user-collected query data with the reference (see Methods).

**Fig. 6.**
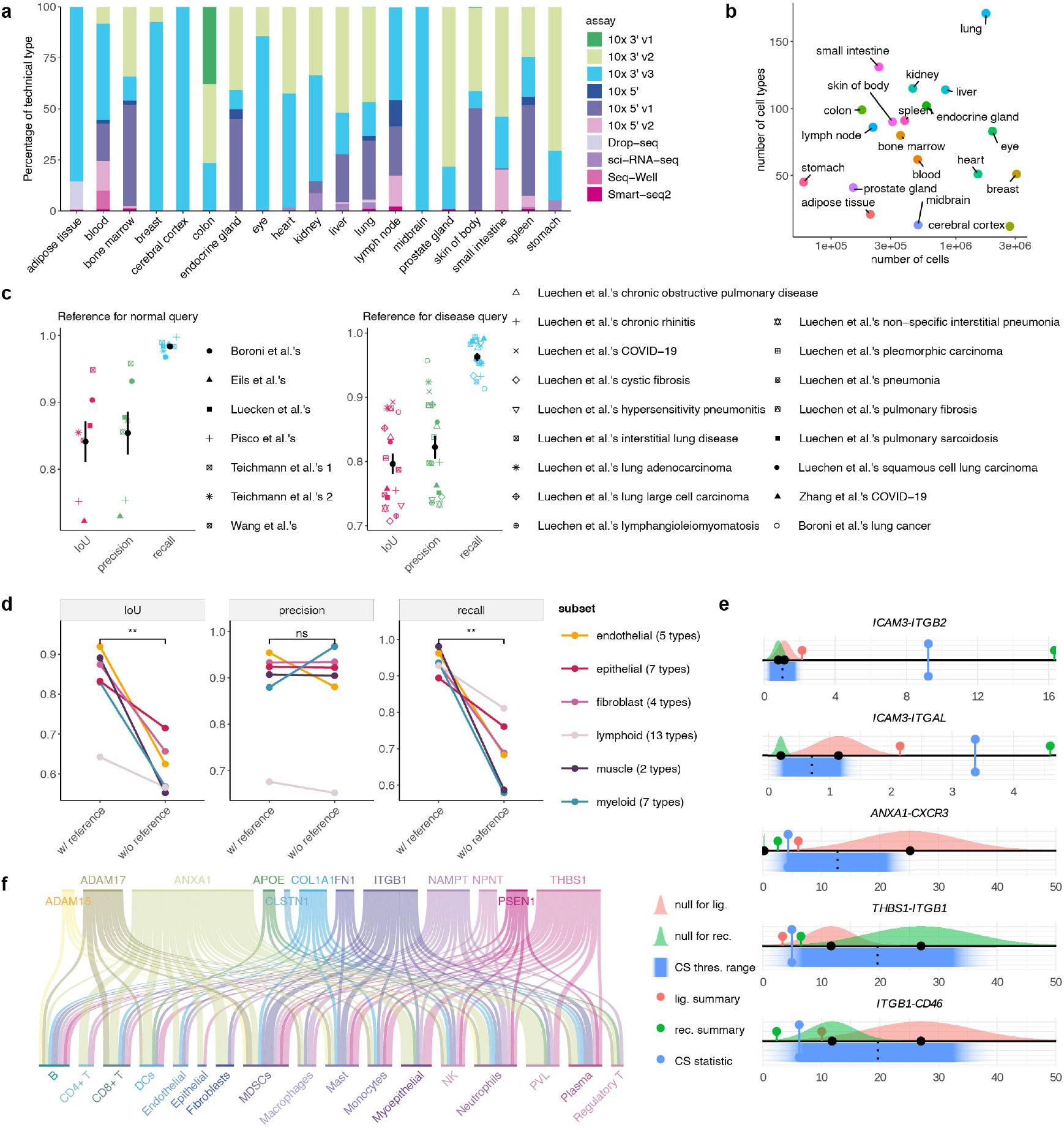
Construction of the first human CCC reference panel and validation of reference-based CCC analysis. **a**. Stacked bar plot showing the proportion of scRNA-seq data from different technical platforms in each tissue CCC reference panel. **b**. Scatter plot showing the number of cells and cell types in each reference panel. **c**. Comparison of results obtained using the reference-based method and conventional CCC analysis in the benchmarking dataset which serves as the ground truth. Each point represents a query dataset, with the mean and standard deviation of each metric annotated. **d**. Pairwise comparison between the results of the reference-based method and conventional CCC analysis using the biased datasets. Each point represents a subset containing a group of similar cell types as the query test. Results from conventional CCC analysis on the entire dataset were used as ground truth. Pairwise t-tests were performed, with ‘ns’ indicating non-significant differences and ‘**’ for *p*-value *<* 0.01. **e**. Examples of communications between cancer epithelial cells and NK cells are demonstrated. Gaussian distributions represent the null expression summaries of ligands and receptors from the reference, with their statistical values mapped onto the query dataset. Blue bands indicate the mapped CS results, with dark blue representing the 95% confidence interval. Correspondingly colored dots show the levels of the respective statistical values in the query data. **f**. Sankey plot showing downregulated CCCs in the query data with cancer epithelial cells as the sender cell type. The first row represents the ligands involved in the downregulated CCCs, while the second row indicates the cell types these ligands act upon via their receptors.

Reference-based CCC analysis using the human CCC reference panel with FastCCC is straightforward, requiring users to provide only the raw gene expression count matrix of their query data. The workflow for reference-based CCC inference implemented in FastCCC involves multiple steps. First, FastCCC preprocesses the query data by converting gene expression to expression ranks in each cell (see Methods), a strategy used in expression quantitative trait mapping studies[38] and single-cell foundation models[39], to minimize batch effects between the query and reference datasets. Second, FastCCC relies on housekeeping genes[40, 41] to adjust for the remaining differences between the two datasets through mapping the reference-based expression summaries onto the query domain. Finally, based on the number of cell types in the query dataset, FastCCC determines the threshold for declaring signficiant and further map the threshold to the query data into a probabilitic distribution to determine the mapped significant threshold for *CS*, thereby identifying significant LRIs in the query data (more details provided in the Methods).

We first validated the use of reference-based CCC analysis using lung data comprising of approximately 1.7 million cells from our constructed human CCC reference panel in two distinct case studies. In the first case study, we constructed three distinct query datasets by extracting three anatomical regions of the lung, including lung parenchyma, the lower lobe of the left lung, bronchus, and the upper lobe of the left lung, while retaining the remaining lung data as the reference. In the second case study, we obtained four normal lung datasets not included in our CCC reference, along with three lung-related disease datasets (one of which encompassed fifteen distinct diseases, each analyzed separately; see Methods). Each of these 21 normal or disease datasets was treated as a query data. For the reference data, we used either the complete lung data from our human CCC reference as the reference panel or the lung reference data from the first case study. As shown in the Fig 6c, the results from reference-based CCC analysis are highly accurate compared to standard CCC analysis using the query data alone. Specifically, across all normal and disease query datasets, the average precision reached 80%, with an IoU of 80% and recall of 95%. In addition, in the second case study, for the four external normal datasets and seventeen disease datasets, the results were nearly identical between the two references(Fig A13 A14), with slight increase in precision and IoU for the larger reference, demonstrating the robustness of reference-based CCC analysis using the constructed human CCC reference.

One important advantage of FastCCC’s reference-based approach is its ability to capture a diverse range of cell types, facilitating the generation of more accurate background distributions and, consequently, more precise CCC analysis. Compared to analyzing the query data alone, incorporating a reference helps mitigate inherent biases in CCC analysis that may arise from the relatively small size or specific experimental conditions of the query data. For example, certain query datasets may involve selective sorting of single cells, where precise isolation of specific cells from a heterogeneous population is conducted, thus artificially inflating the mean *CS* for all LRI pairs associated with these cell types in the null distribution. Such inflation of null statistics can result in a loss of power and failure to detect potentially up-regulated interactions. By leveraging our comprehensive human CCC reference, which encompasses a diverse range of cell types, such biases can be effectively mitigated.

To illustrate this benefit, we used the dataset from Teichmann et al., which contains a total of ∽318k cells across 38 cell subtypes that belong to six major cell type groups: endothelial, epithelial, fibroblast, lymphoid, myeloid, and muscle. We conducted conventional CCC analysis in the full dataset and treated the results as the ground truth. In addition, we split this dataset into six subsets based on the major cell type groups and conducted independent CCC analyses for each subset, which exhibited pronounced imbalances in cell type proportions compared to the full dataset. This imbalance significantly affected the baseline levels of *CS* statistics for certain LRI sets associated with specific cell types, potentially introducing biases when applying CCC analysis directly to the subdivided datasets rather than the full dataset. Ideally, an accurate inference method should yield consistent CCC results regardless of the cell types present in the dataset, leading to similar results as inferred from the full dataset. However, in some extreme cases – such as myeloid and muscle major group – the IoU and recall from CCC analysis using the query data alone dropped to below 60% when compared to the results of the full dataset used as ground truth (Fig 6d). In contrast, inference using the human CCC reference panel consistently outperformed inference using query data alone across all analyses. Specifically, using the lung CCC reference panel achieved an average consistency exceeding 80% compared to the ground truth obtained from the full dataset. For all subdivided datasets, recall consistently surpassed 90%, while precision remained at the same level. Even in extreme scenarios – where the query dataset contained only a single cell type and conventional CCC methods failed to produce any autocrine interaction results, reference-based CCC analysis using the human CCC reference panel maintained an average IoU exceeding 80% for single-cell-type query datasets (Fig A15).

To further demonstrate the benefits of FastCCC’s reference-based approach, we utilized the breast reference from our human CCC reference panel, comprising over 3 million cells and 51 cell types from 323 healthy samples, to analyze an independent breast tumor dataset[42] obtained externally. Despite the differences in the resolution and ontology of cell type annotations between the reference and query datasets, FastCCC’s versatile reference-based approach enables flexible merging of subtypes or mapping of cell type labels to align the meta-information of the reference and query datasets. In the reference-based analysis, we found that significant LRIs in breast cancer were predominantly observed between specific immune and stromal cell types and cancer epithelial cells. For instance, cancer epithelial cells, as senders, influence immune and stromal cells through receptors such as TIGIT, ITGB1, PIK3CB, and a range of chemokines and cytokines (Fig A16)—findings consistent with previous research[43– 48]. These receptors are involved in pathways extensively linked to cancer progression, including cell adhesion molecules, whose disruption is a hallmark of cancer [44, 45]; to PI3K-Akt signaling, a key regulator of survival, metastasis, and metabolism [46, 47]; and to ECM-receptor interaction, crucial in breast cancer development [48] (Other top enriched KEGG pathways see Fig A17). Compared to conventional CCC analysis using large-scale normal breast tissue samples, reference-based CCC analysis enables precise identification of contributions to significant *CS* statistics. For example, in LRIs between cancer epithelial and NK cells, such as ICAM3-ITGB2 and ICAM3-ITGAL, both ligands and receptors are expressed at significantly higher levels in the query dataset than in normal conditions(Fig 6e). In these cases, *CS* in the query dataset are over ten times the standard deviation beyond the *CS* threshold from the reference, indicating a strong communication between these two cell clusters.

The second important advantage of FastCCC’s reference-based analysis lies in its ability to detect reduced CCC levels in the query data compared to the reference, which is composed of normal tissues. Specifically, conventional methods focus only on identifying significantly enhanced CCCs exceeding certainly threshold. In contrast, FastCCC leverages a CCC reference panel constructed from normal tissues to perform additional analysis, enabling the identification of CCCs with significantly reduced interactions relative to the reference, in addition to significantly enhanced CCCs. For instance, in ANXA1-CXCR3, THBS1-ITGB1, and ITGB1-GD46 interactions between cancer epithelial and NK cells(Fig 6e), ligand expression in cancer cells is significantly lower in the query dataset compared to the reference, regardless of receptor expression levels. The overall *CS* values fall more than two standard deviations below the normal reference threshold, suggesting markedly reduced communication levels.

Notably, across the query dataset, 12 cancer epithelial ligands, including ANXA1, THBS1, ITGB1, and ADAM17, were consistently associated with all other immune cells through LRIs that fell below 1.96 times the standard deviation of the normal reference threshold (Fig 6f). Importantly, literature extensively links these ligands’ overexpression with increased tumor growth, enhanced tumor cell migration, aggressiveness, and poor prognosis[49–56]. To further validate the clinical relevance of FastCCC results, we separately analyzed single-cell data from triple-negative breast cancer (TNBC) patients—a more aggressive breast cancer subtype with a worse prognosis—accounting for less than 30% of the total dataset. Reference-based CCC analysis in TNBC samples revealed notable differences: LRIs associated with most of the aforementioned ligands, such as ANXA1, THBS1, and NAMPT, were no longer significantly downregulated (Fig A19). Additionally, ITGB1 and ADAM17-related LRIs showed a dramatic reduction in the number of significantly decreased interactions, involving only 2–4 CCCs. For these remaining downregulated LRIs, the reduction primarily resulted from decreased receptor expression.

These findings demonstrate that reference-based comparisons effectively provide background distribution information, closely approximating the results of direct CCC inference methods applied to datasets with large, diverse cell populations, aligns with clinical phenotypes and offers mechanistic insights into tumor behavior and microenvironmental changes. This reference-based approach effectively mitigates biases associated with small datasets and provides more comprehensive characterization of CCCs, enabling researchers to obtain a more accurate representation of CCCs and facilitating a deeper understanding of intercellular communication.

## 3 Discussion

FastCCC addresses critical limitations in CCC analysis by introducing a highly scalable, permutation-free framework that integrates FFT-based *p*-value computation, a modular scoring system, and reference-based analysis. These innovations enable scalable, robust, and effective analysis of large-scale datasets, significantly surpassing the capabilities of existing tools.

In diverse biological contexts, FastCCC reliably identifies meaningful CCCs. Notably, it captured interactions between endothelial and immune cells mediated by the C3 ligand and its receptors, correlating with COVID-19 severity, and revealed dynamic LRIs during T-cell development in thymic tissue, as well as the correlation between changes of down-regulated LRIs with clinical phenotype in breast cancer. These findings highlight its potential to uncover critical pathways in complex tissues and disease states. The incorporation of a comprehensive human CCC reference panel further enhances analytical power, enabling systematic comparisons and facilitating the discovery of conserved and context-specific interactions. Beside those advantages, one potential limitations the reference constructed by FastCCC contains human samples only and are not yet applicable other species. Extending FastCCC to non-human datasets for reference-based analysis would further enable the exploration of CCCs in diverse model organisms, broadening its applicability to comparative and evolutionary studies.

In summary, FastCCC represents a transformative advance in single-cell CCC analysis, combining efficiency, scalability, and biological insight. Its flexibility and reference-based capabilities pave the way for more comprehensive studies of intercellular communication across health and disease.

## 4 Methods

### 4.1 A general CCC testing framework

We begin with a candidate list of ligand-receptor pairs, which can be either user-defined or obtained from existing databases. For each ligand *l* and receptor *r*, we analyze one pair of cell types at a time. For each such quadruplet, we compute an communication score (*CS*) to quantify the coordinated expression of ligand *l* in cell type *c*_*a*_ and receptor *r* in cell type *c*_*b*_ . The *CS* score is represented in general form as follows:

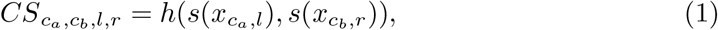

where 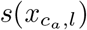 is a scalar and represents the gene expression summary of ligand *l* across cells of cell type 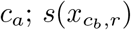 is also a scalar and represents the gene expression summary of receptor *r* across cells of cell type *c*_*b*_; and *h*(·, ·) is a function that measures the coordinated expression between 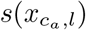 and 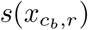.

The *CS* statistic described in equation 1 is quite versatile, accommodating various functional forms for *h*(·) and *s*(·). For *s*(·), several summary statistics can be used to provide a point summary of ligand/receptor gene expression levels across cells in a given cell type. In the default version of FastCCC, we consider four such statistics: mean, median, 3^rd^ quartile, and 90^th^ percentile. These statistics effectively captures diverse cell-cell interaction patterns; for example, the mean expression reflects average expression across cells, capturing interactions that are prevalent in a large fraction of cells, while the 90^th^ percentile is effective in capturing interactions among a small subset of cells with high expression levels. Importantly, these statistics are also applicable when the ligand or receptor is a molecular complex comprising multiple subunits. In such case, FastCCC computes the point summary for each subunit separately and then considers either the minimum or average expression across the subunits as the final *s*(·). The minimum expression is effective in scenarios where all subunits must be expressed for the ligand or receptor complex to function, whereas the average expression among all subunits offers a more lenient assessment that captures the overall activity across the components. Again, while the default version of FastCCC offers minimum and average as two options, its framework is general and can easily incorporate additional alternatives.

For *h*(·), we consider two distinct functional forms. The first form is the arithmetic average of the ligand and receptor expression summaries, represented as

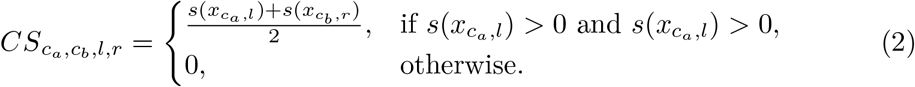

The second form is the geometric average of the ligand and receptor expression summaries, represented as

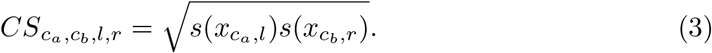

The arithmetic average is particularly effective in capturing the central tendency of the data and in detecting strong interactions where either the ligand or receptor is highly expressed. In contrast, the geometric average reflects interactions that may scale multiplicatively, providing a better representation of ligand-receptor levels that span several orders of magnitude.

Thanks to the general framework of FastCCC, it includes many previously proposed *CS* statistics for CCC as special cases. For example, when 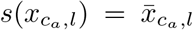 and 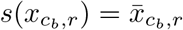 are defined as the average expression measurement of ligand and receptor, respectively, and *h*(·) follows the same format as equation 2, representing the average of these expression measurements, then the *CS* in equation 1 simplifies to the test statistics used in CellPhoneDB. Importantly, the statistical power to detect CCC inevitably depends on how well these chosen functions of *h*(·) and *s*(·) capture the true interaction pattern between ligand and receptor for specific cell type pairs. Unfortunately, the true underlying CCC patterns for any ligand-receptor and cell type pair are unknown and may vary considerably across quadruplets. To ensure robust identification of CCC across various scenarios, we incorporated two distinct *h*(·) functions, four *s*(·) functions for single-unit ligand or receptor, and 2 *s*(·) aggregation functions for ligand or receptor complex. This combination results in a total of 16 scoring methods implemented in FastCCC (detailed below and also Table A3). We fit our model for each combination of *h*(·), *s*(·), and aggregation function, calculate the corresponding *p*-value, and combine these 16 *p*-values through the Cauchy *p*-value combination rule to yield a single *p*-value as the final evidence for interaction. To apply the Cauchy combination rule, we convert each *p*-value into a Cauchy statistic and aggregate all Cauchy statistics through summation in the form of 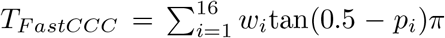,where we used equal weights of *w*_*i*_ = 1*/*16. Afterwards, we convert the summation back to a single *p*-value based on the standard Cauchy distribution in the form of *p*_*F astCCC*_ = 1*/*2 −tan^−1^(*T*_*F astCCC*_)*/π*. The Cauchy rule takes advantage of the fact that combination of Cauchy random variables also follows a Cauchy distribution regardless of whether these random variables are correlated or not. Therefore, the Cauchy combination rule allows us to combine multiple potentially correlated *p*-values into a single *p*-value without loss of type I error control. This approach allows for flexibility in capturing different patterns of interaction based on varying expression metrics and interaction models.

### 4.2 Detailed choices of *s*(·)

We provide the detailed functional forms for *s*(·) in this section. For a single-unit ligand or receptor, we consider four different choices for *s*(·). The first choice is the mean, represented as

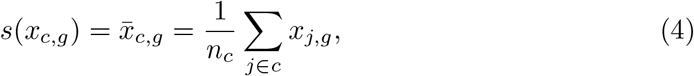

where the subscript *g* ∈ {*l, r*} represents a single unit of either ligand and receptor; the subscript *c* ∈ {*c*_*a*_, *c*_*b*_} represents either cell type *c*_*a*_ or *c*_*b*_; *n*_*c*_ represents the number of cells belonging to cell type c; *x*_*j*,*g*_ represents the expression level of ligand or receptor for the j^th^ cell in cell type c; and 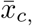.represents the average expression level of ligand or receptor for cell type c.

The second to fourth choices for *s*(·) are three order statistics of gene expression, represented as the median, 3^rd^ quartile and 90^th^ percentile from an arbitrary expression distribution. Specifically, for a quantile value *q* (*q*=50, 75, or 90), we compute the *k*^th^ order statistic in the form of

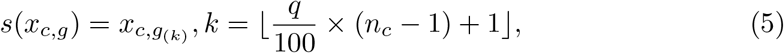

where 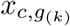 represents the *k*^th^ value in the sorted gene expression levels of the ligand or receptor within the corresponding cell type, arranged in ascending order; the value of *k* is rounded down to determine the rank position to be calculated.

In the setting when either the ligand or receptor is in the form of a multi-subunit complex, the expression summary statistics for the complex is aggregated by taking either the average or the minimum of its subunits. Let *n*_*g*_ represent the number of subunits; we have the following two choices to compute the final *s*(·):

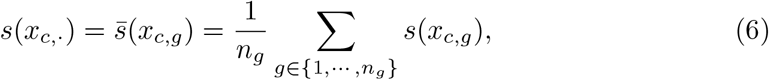

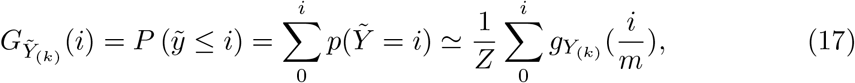

### 4.3 Convolution-based *p*-value calculation for *CS* statistics

After obtaining the above *CS* statistics, the next step is to test whether the observed *CS* statistic is as extreme as, or more extreme than, what would be expected under the null hypothesis. Let *n*_*a*_ denote the number of cells in cell type *c*_*a*_, and *n*_*b*_ the number of cells in cell type *c*_*b*_. The null hypothesis is defined based on two cell types with the same sample sizes *n*_*a*_ and *n*_*b*_, but with ligand and receptor expression randomly drawn from the entire distribution across all cells. The standard approach for computing the corresponding *p*-value involves randomly sampling *n*_*a*_ and *n*_*b*_ cells from the population, which is achieved practically by permutating cell type labels or shuffling the cell type identities, and then recalculating the *CS* statistic. This permutation process is repeated at least thousands of times to construct a null distribution of *CS* statistics. The proportion of *CS* statistics from the permuted null distribution that exceeds the observed *CS* gives the *p*-value. Unfortunately, this permutation-based *p*-value computation procedure is computationally intensive and can be sensitive to the number of permutations employed.

Here, we take an alternative approach to obtain the distribution of *CS* statistics under the null using scalable analytic solutions rather than permutation-based numeric solutions. Specifically, we first denote *f*_*h*_(·) as the probability density function (PDF) for *CS* statistics under the null. Our objective is to compute this PDF, which allows us to further compute the *p*-value corresponding to an observed *CS* statistic: 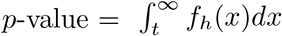 for an observed *CS* statistic equal to t. To achieve this, we derive our key insights from the fact that *f*_*h*_(·) function can be effectively expressed as a convolution of two separate functions. To see this, we first recognize that the first *h*(·) function defined in equation 2 represents the average of the summary levels of ligand and receptor, while the second *h*(·) function defined in equation 3 is similarly the average of the two, but on the log scale. Therefore, for an *h*(·) function defined in equation 2, we have

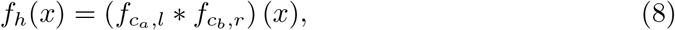

where 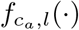 denotes the PDF describing the distribution of the summary expression level of ligand in cell type 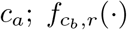 denotes the PDF describing the distribution of the summary expression level of receptor in cell type *c*_*b*_; and *f* ∗*f* denotes the convolution of two functions, defined as

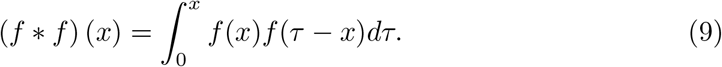

For an *h*(·) function defined in equation 3, we similarly have

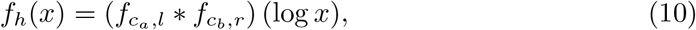

where 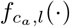 denotes the PDF describing the distribution of the log-scale summary expression level of ligand in cell type *c*_*a*_; and 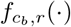 denotes the PDF describing the distribution of the log-scale summary expression level of receptor in cell type *c*_*b*_.

Using the above insights, we first obtain the two PDF functions 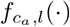 and 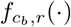 separately. These functions depend on the specific choice of *s*(·) and the presence or absence of a multi-subunit complex, as detailed in later sections. Afterwards, we compute the PDF *f*_*h*_(·) based on their convolution. In the simplest case where *f*_*h*_(·) function defined in equation 2 represents the average of the summary levels of ligand and receptor and where none of the two cell types is rare, this *f*_*h*_(·) function can be directly derived based on asymptotic properties and is in the form of a normal distribution. In the more general case, we utilize fast Fourier transformation (FFT) for computation, accommodating any arbitrary distributional forms and ensuring scalable computation.

### 4.4 Probability density functions for single-unit ligand or receptor with different choices of *s*(·)

Here, we provide details for obtaining the probability density function *f*_*s*_(*x*)(*s* = *c*_*a*_, *l* or *c*_*b*_, *r*) for the summary expression level of a single-unit ligand or receptor in a specific cell type. To simplify presentation, we focus on the distribution of the summary expression level of *x* rather than the log-scale summary expression level of log *x*, as the derivations and resulting forms are largely similar.

First, we consider the case where the *s*(·) function represents the expression mean as described in equation 4. For non-rare cell types where the number of cells exceeds 30, we apply the central limit theorem (CLT) to obtain the sampling distribution of the mean as an asymptotical normal distribution. Such normal distribution has mean *µ*_*g*_ and variance 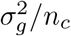,where *µ*_*g*_ and 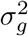 are the mean and variance of gene expression in all cells regardless of the cell type (*g* ∈ {*l, r*}), respectively, and *n*_*c*_ represents the number of cells in the cell type of focus (*c* ∈{*c*_*a*_, *c*_*b*_}). For rare cell types where the number of cells does not exceed 30, we cannot apply CLT. Instead, we use convolution to calculate the exact distribution of *f*_*s*_(*x*). Because observing a mean value of 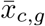 is equivalent to observing the corresponding total expression value of 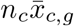,we have 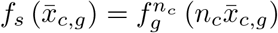,where 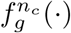 is defined as a convolution across *n*_*c*_ cells in the form of

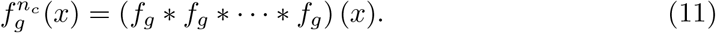

where *f*_*g*_ represents the PDF function of expression level for each cell; and ∗denotes convolution operation defined in equation 9 earlier. We compute such convolution based on fast Fourier transformation (FFT) on any arbitrary distribution *f*_*g*_. Specifically, we apply log-transformed normalization to the expression profiles, discretize *f*_*g*_ by partitioning the domain into evenly spaced segments with a minimum precision of 0.01, count the frequency of each segment to capture the discrete probability of expression within each segment, and apply FFT to perform convolution and calculate the PDF of *f*_*s*_(*x*).

Next, we consider the case where the *s*(·) function represents the median, 3^rd^ quartile, or 90^th^ percentile of the expression level. Here, we denote *q* as the percentile of interest (*q*=50, 75, or 90) and obtain the corresponding *k*^th^ order statistic, where *k* = ⌊*q/*100× (*n*_*c*_ −1) + 1⌋ . We aim to obtain the PDF for *k*^th^ order statistic from an arbitrary discrete distribution, regardless of whether the cell type is rare or common. To do so, we denote *X*_1_, *X*_2_, · · ·, *X*_*n*_ as *n* observed expression values for the ligand or receptor in the cell type of focus. We sort them such that *X*_(1)_ *< X*_(2)_ *<* · · · *< X*_(*n*)_.

We denote *f* (*x*) as its PDF. The distribution for *k*^th^ order statistic from the expression level is:

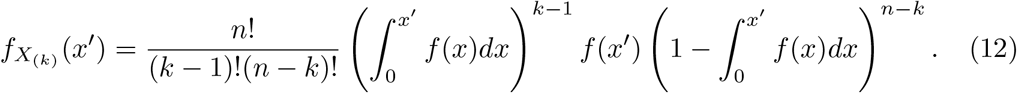

To infer such distribution, we create a reference sample, denoted as *Y*_1_, *Y*_2_, ···, *Y*_*n*_, which are *n* samples randomly drawn from a uniform distribution *U* (0, 1). We also sort them such that *Y*_(1)_ *< Y*_(2)_ *<* ···*< Y*_(*n*)_. The distribution for *k*^th^ order statistic from this reference distribution can be derived as follows

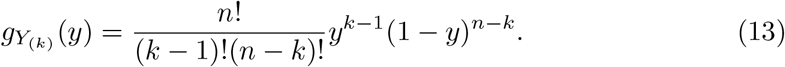

We connect the expression data to the reference distribution through the cumulative distribution functions (CDF) of *X* and *Y* . Specifically, we denote *F*_*X*_ and *G*_*Y*_ as the CDF of *X* and *Y*, respectively, with range at [0, 1]. We identify the map function between the two as *F*_*X*_ (*x*) = *G*_*Y*_ (*y*). It is straightforward to deduce that

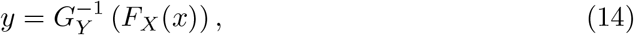

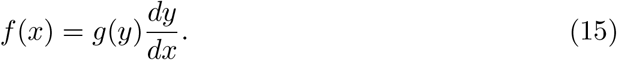

Therefore, we have:

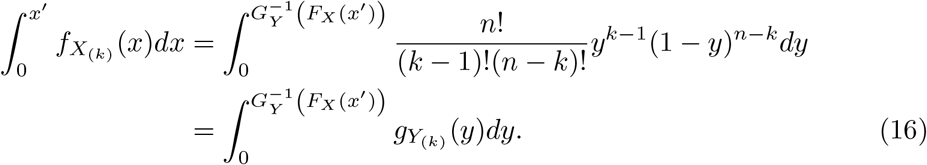

With the above equation, we obtain the cumulative distribution for the *k*^th^ order statistic for a set of uniform random variables, and map *x* to *y* through CDF of the gene expression level to further obtain the cumulative distribution of the PDF for *X*_(*k*)_.

To facilitate computation, we discretize the interval [0, 1] evenly into *m*=100,000 segments, enabling us to compute the CDF for the *k*^th^ order statistic of a set of uniform random variables. This approach allows us to further calculate the CDF and subsequently the PDF for the summary expression level. In practice, for *i* = 1, 2, · · ·, *m*, we compute 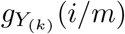.For a random variable 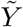 with a discrete uniform distribution over the possible values *i* = 1, 2, · · ·, *m*, the approximate solution for the CDF of the *k*^th^ order statistic of 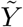 is given by:

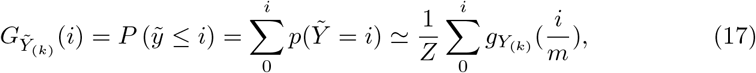

where *Z* is the normalization constant used to ensure that the resulting distribution integrates to 1, maintaining the properties of a valid probability distribution. Thus, we can calculate the probability mass function 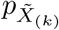 for the *k*^th^ order statistic of any discrete gene expression random variable 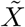,obtained from the data, with all possible values *x*_1_, *x*_2_, · · ·, *x*_*l*_ (*x*_1_ *< x*_2_ *<* · · · *< x*_*l*_). Specifically, we have:

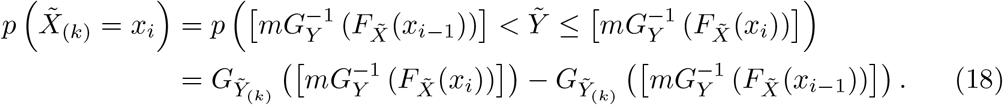

### 4.5 Probability density functions for ligands or receptors with multiple subunits

For ligands or receptors with multiple subunits, we first consider the case where the expression summary is the average of *s*(·) for all subunits as defined in equation 6. In the simplest case where the expression summary for each subunit follows a normal distribution, the distribution for the average is also a normal distribution: the mean equals the average of the means of each subunit and the variance is 1*/n*^2^ times the sum of variances of each subunit. In the general case where the expression summary of each subunit follows an arbitrary distribution, the distribution for the average is equivalent to the convolution of *n*_*g*_ discrete distributions in the form of

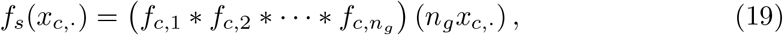

where *f*_*c*,*g*_(·) denotes the PDF for each subunit *g* (*g* ∈ {1, · · ·, *n*_*g*_}) in cell type *c*.

Next, we consider the case where the expression summary is the minimum of *s*(·) across subunits as defined in equation 7. In this case, we denote *F*_*c*,*g*_ to represent the CDF of 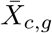 and CDF *F*_*s*_ for the minimum expression summary variable. We obtain the CDF for the minimum of *n*_*g*_ random variables from the *n*_*g*_ subunits as

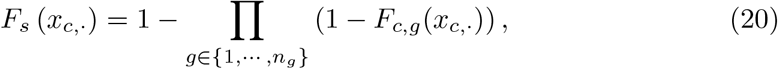

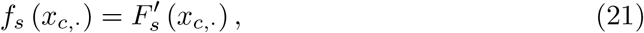

where *f*_*s*_ (*x*_*c*,·_) denotes the corresponding PDF function.

### 4.6 Valid candidate LRIs

Before running FastCCC, we perform ligand-receptor pair filtering to focus on a set of valid candidate LRIs for interaction analysis. To do so, we follow existing approaches to filter out genes with a large amount of zero expression in certain cell types when considering them as candidate ligands or receptors. In FastCCC, we use CellPhoneDB’s default threshold of 10%, meaning that if a gene’s expression is zero in more than 90% of cells within a given cell type, we no longer consider LRIs involved by that gene for that cell type, and its p-value is set to 1. These LRIs that are involved in the actual computation are referred to as “valid LRIs,” as mentioned in the main section.

All of our evaluation results are based on valid LRIs. Additionally, FastCCC provides an option to follow the recommendations of difference-assembly-based tools and use our probability distribution toolkit to directly screen for differentially expressed genes (DEGs). Specifically, FastCCC can be used to estimate the null distribution of gene expression within each cell type cluster, thus directly calculating p-values using observed gene expression without relying on external tools to identify DEGs. This allows FastCCC to efficiently pinpoint DEGs across specific cell types and further incorporate DEGs as an additional filter for candidate LRIs to ensure a rigorous selection process. In particular, such filter requires both ligands and receptors to be highly expressed in their respective cell clusters, thereby reducing false signals. Users have the option to choose whether to include DEGs as a selection criterion based on their specific analysis needs. This design offers flexibility, allowing researchers to adapt the toolkit to a wide range of biological and computational scenarios by freely combining different operational modules in a heteromer modeling framework and analytically deriving the null distribution of the scores.

### 4.7 Reference-based CCC Inference

Reference-based CCC analysis in FastCCC requires both query data and reference data. The query data, typically a smaller user-collected dataset, is provided as a raw gene expression count matrix. The reference data, significantly larger in scale, is provided and processed through FastCCC. The reference data includes essential information such as the reference tissue name, LRIs DB used, the types and numbers of cell types included, the PDFs of gene expression, and some precomputed information necessary for the FastCCC inference workflow. No reference expression profile data or meta info is stored or used in the reference panel, ensuring scalability and conciseness in data representation. To support reference-based CCC analysis, we have constructed a human CCC reference panel, with details provided in the next section.

For both query and reference data, FastCCC performs basic quality control to filter out cells with abnormally low gene expression levels (i.e. cells with *<* 50 expressed genes). Afterwards, FastCCC adapts rank-based batch correction and data normalization techniques to remove potential batch effects between query and reference data. Specifically, in each dataset, it sorts the non-zero expressed genes in ascending order, divides them evenly into *n*_bin_ predefined intervals, and assigns each gene a rank based on its interval. The smallest non-zero expression value was assigned a rank of 1 and the maximum value was set to *n*_bin_. Genes with zero expression remained as 0.

While the rank-based batch correction procedure is effective, the normalized query and reference data inevitably retain some remaining level of noise. To further improve batch correction, we utilize housekeeping genes[40] to estimate the remaining noise and directly align the two datasets. To do so, we assume that the mean expression of each housekeeping gene *g* in the reference (*X*_*r*,*g*_) and query (*X*_*q*,*g*_) datasets satisfies

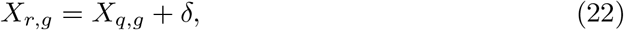

where *δ* follows a normal distribution with a mean of 0 and a standard deviation proportional to *X*_*r*,*g*_:

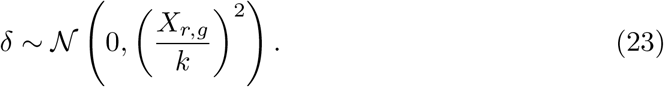

We estimate the parameter *k* using all the housekeeping gene means, which are recorded during the reference construction step, and the corresponding values in the query data. *k* is set to a minimum value (default set to 3) if the estimate value falls below it, mitigating excessive uncertainty.

Once the noise parameter *k* is estimated, all statistics in the reference data can be mapped to the query domain using properties of a Gaussian distribution. Specifically, a point statistic in the reference, *X*_*ref*_, is mapped to a corresponding value in the query data, denoted as *X*_*query*_, as follows:

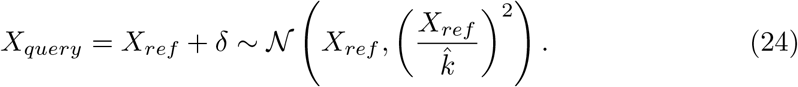

Importantly, the above equation not only gives out the mapped value *X*_*query*_ but also provides a confidence interval for the mapped value, allowing us to readily assess uncertainty.

With the above setup, we calculate the *CS* statistics for each LRI in the query dataset using equations 1, 2, 4, and 6, along with the corresponding ligand and receptor expression summary values *s*(·), with a precision of 0.01. In parallel, we use the null distribution of expression summary of ligands and receptors obtained from the reference data to calculate a *CS* significance threshold corresponding to a *p*-value of 0.05. This threshold in the reference data is mapped to the query data using equation 24 to yield both the query data threshold and the 95% confidence interval associated with this mapped threshold. Afterwards, we use the confidence interval to compute the lower and upper bounds associated with the mapped query data threshold. Additionally, we also obtained the null expression summaries of ligand and receptor from reference and mapped them using equation 24. Based on the relationship between the *CS* values of LRIs in the query dataset and the corresponding mapped *CS* threshold confidence interval, we considered the following five scenarios to determine significance. 1. LRIs with *CS* values above the upper bound are considered as significantly communicate. 2. those with *CS* values below the lower bound are deemed insignificant. For LRIs with CS values within the threshold interval, we conducted further comparison with the reference to determine significance. First, we check whether the corresponding sender-receiver’s LRI is significant in the reference. 3. If both the sender and receiver cell types exist and the LRI is significant in the reference, the ligand and receptor’s expression summaries in the query dataset are compared with the lower bound of the mapped corresponding null summaries from reference. And if both intensity exceed the lower bound, the LRI is considered significant. 4. On the other hand, if the LRI is not significant in the reference and the ligand and receptor’s expression summaries in the query dataset do not exceed the upper bound, the LRI is considered insignificant. 5. In cases where the conditions above do not apply – such as when *CS* falls within the mapped threshold interval while the sender or receiver cell type is absent from the reference, or if ligand and receptor expression comparisons with the reference yield conflicting results (e.g., one increases while the other decreases, leading to a CS value near the threshold) – the significance of the LRI is assessed using query data alone by comparing the *CS* value to its null distribution.

Through the above process, FastCCC enables reference-based CCC analysis by comparing the query and reference data, enabling the identification of CCC signals that conventional CCC analysis might overlook. Reference-based CCC analysis is particularly valuable in scenarios where such CCCs are systematic and global, rendering them challenging to detect when using the query dataset alone as the background.

### 4.8 Construction of Human CCC Reference

To facilitate reference-based CCC analysis with FastCCC, we constructed a human CCC reference panel using the CellxGene[37] platform (version dated 2024-07-01). Specifically, we first obtained 19 single-cell RNA sequencing datasets from the platform, each representing a different normal human tissue, with each dataset containing at least 150,000 cells (an exception is stomach tissue, which contains over 60,000 cells and is included as a reference due to its frequent use in studies). In the process, we selected “Homo sapiens” as the species, “normal” as the disease condition, set “is primary data” to false to use prefiltered data, and used the “tissue general” option to select datasets for the following tissues: brain, breast, eye, lung, heart, liver, endocrine gland (or thymus), blood, kidney, respiratory system, spleen, bone marrow, skin of body, small intestine, lymph node, adipose tissue, colon, nose, and prostate gland. For brain tissues, we further refined the selection using the “tissue” option to include two representative regions: midbrain and cerebral cortex. Detailed information about the reference data for these different tissues is provided in the Supplementary information. All datasets used HGNC symbols as gene names, which were retained during processing. For cell type information, we followed the CellxGene platform guidelines, using the cell type labels provided by the platform according to the Cell Type Ontology standard as input for FastCCC. Additionally, we employed the LRIs database provided by CellPhoneDB v5[8] as the candidate list for constructing the reference. For each dataset in turn, we conducted rank-based batch correction and normalization desribed in the previous section. We then obtain PDFs and summary information in the data in the form of mean of the ranked gene expression. We only retained the necessary information required by FastCCC for reference construction as detailed in the previous section to ensure scalability and conciseness in the reference. For example, raw count matrices and other dataset-specific information were neither saved nor used during reference-based CCC analysis.

We used the lung dataset from the constructed human CCC reference panel to illustrate reference-based CCC analysis in two distinct case studies. In the first case study, we extracted lobe-related and other positions from the dataset to serve as query, and used the remaining lung data as the reference (which contains over 860k cells). In the second case study, we downloaded supplementary lung normal datasets excluded from our CCC reference panel to serve as the query data, mimicking the smaller size typical of user-collected dataset. We ensured no cell overlap between the reference and query data by using observation joinid, a unique identifier assigned to each cell, which is required when uploading data to the CellxGene platform.

### 4.9 Simulations and benchmarking: data sources, designs, and evaluations

#### Data sources

We first conducted comprehensive simulations and benchmarking using eleven datasets, covering distinct biological contexts and scenarios. These datasets encompass cell numbers ranging from several thousand to two million, with the number of cell types varying from under five to over 50, thereby covering the majority of scenarios encountered in practical applications. The datasets include:

1. CellPhoneDB tutorial dataset, designated as CPDBTD. This dataset is an example dataset provided by CellPhoneDB, used to validate the correctness of our algorithm’s theoretical framework and implementation. It contains 3,312 cells across 40 cell types. Together with CellPhoneDB and a modified version based on the *k*-th order statistic, this dataset was used to verify equations (8–21).
2. Human Ileum dataset. This dataset is part of a scRNA-seq study measuring human intestine[57], especially the ileum segmentation. It contains 16,819 genes and 5,980 cells across 7 cell types.
3. Human Fibroblast dataset. This dataset is part of an integrated scRNA-seq analysis of the normal gastrointestinal (GI) tract, esophagus and stomach measured by 10X Genomics[58]. It contains 33,234 genes and 5,754 cells across 3 cell types.
4. Human ECGE dataset. This dataset collects snRNA-seq data measured in human entorhinal cortex (EC) and ganglionic eminence (GE) germinal zones[59]. It contains 21,563 genes and 7,618 cells across 10 cell types.
5. Human Breast dataset, designated as BreastB, is a subset of data from the Human Breast Cell Atlas (HBCA) study[60]. This dataset includes B cells and comprises 33,234 genes and 12,510 cells across 5 cell types.
6. Human HCA Kidney dataset, designated as Kidney. This dataset contains scRNA-seq data from normal regions of kidney tumor-nephrectomy samples collected from 14 individuals[61]. It contains 37,073 genes and 48,783 cells across 28 cell types.
7. Human AnG dataset, designated as Angular. This dataset is part of an snRNA-seq study using SMART-seqv4 RNA-sequencing that measures eight cortical areas in the human brain[62]. It focuses on the angular gyrus (AnG) area and includes 2,0981 genes and 110,752 cells across 18 cell types.
8. Human Fetal Retina dataset, designated as Retina. This dataset comprises single-nucleus multiome data from the developing human retina, obtained from 12 donors using 10X Chromium Single-Cell Multiome ATAC + RNA-seq[63]. Our study uses only snRNA-seq data, which includes 36,503 genes and 205,619 cells across 8 cell types.
9. Human GBmap dataset. This dataset consists solely of scRNA-seq data obtained via 10X Genomics, covering a large-scale measurement of 240 patients diagnosed with glioblastoma[64]. The dataset consists of 28,045 genes and 338,564 cells across 17 cell types.
10. Human AIDA (Asian Immune Diversity Atlas) dataset. This dataset includes scRNA-seq data obtained via 10X Genomics, encompassing 503 healthy donors from seven population groups across five countries[65]. It contains 36,266 genes and 1,058,909 cells across 33 cell types. A subset version was also used for running Cell-PhoneDB. For this analysis, we focused on 2,000 highly variable genes (HVGs), designated as AIDA hvg.
11. Human HLCA (integrated Human Lung Cell Atlas) dataset. This dataset integrates 49 studies collected from the human respiratory system into a large-scale atlas data[66]. It consists of 56,295 genes and 2,282,447 cells across 51 cell types. Similarly, CellPhoneDB was applied to 2,000 HVGs, designated as HLCA hvg.

Besides the above 11 datasets, we also used another 11 datasets for benchmarking our human CCC reference panel. The datasets include:

1. Disease samples from MDCs altas : This dataset integrates 73,872 cells and 11 cell types. It is a partial dataset from the multi-tissue myeloid-derived cells single-cell atlas of tumor and healthy samples developed by Boroni et al.[67] We filtered cells from lung tissue with a lung cancer phenotype for testing, and this dataset is therefore referred to as Boroni et al.’s lung cancer dataset.
2. Disease samples from COVID-19 dataset: This dataset is the same one used in our case study when we applied FastCCC to large-scale COVID-19 data[18]. We only used data from the lungs of critically ill patients, containing 42,757 cells and 19 cell types. We designated this data as Zhang et al.’s COVID-19.
3. Disease samples from Human HLCA dataset: This dataset consists of single-cell RNA sequencing data from various lung-related diseases in the HLCA[66], including pulmonary fibrosis, lung adenocarcinoma, and 13 other diseases. Each disease is treated as a separate query dataset. For consistency in naming, in the reference bench-marking study, we used the format “Author + Disease Name,” for example, “Luechen et al.’s cystic fibrosis.”
4. Normal samples from MDCs atlas: This dataset contains 87,370 cells and 11 cell types. It is a partial dataset from the MDCs atlas mentioned earlier. We filtered cells from normal lung tissue for testing, and we referred to this data as Boroni et al.’s dataset.
5. Normal samples from LungMAP: This dataset is a large-scale snRNA-seq dataset of the human lung from healthy donors of approximately 30 weeks, 3 years, and 30 years of age[68]. We used the healthy samples as the query for testing, and it contains 46,500 cells and 27 cell types. It is referred to as Wang et al.’s dataset.
6. Normal lung samples from HCA: This dataset consists of 39,778 cells and 9 cell types derived from normal lung tissue[69]. It is referred to as Eils et al.’s dataset.
7. Normal samples from Human HLCA dataset: This dataset comprises the normal samples from the HLCA, containing over 333,468 cells. It is designated as Luechen et al.’s dataset.
8. Normal lung samples from Tabula Sapiens: Tabula Sapiens is a benchmark first-draft human cell atlas comprising over 1.1 million cells from 28 organs of 24 normal human subjects [70]. For our analysis, we used the normal lung samples from this atlas, containing 65,847 cells and 34 cell types, referred to as Pisco et al.’s dataset.
9. Normal lung samples from CellHint: This dataset, organized by Teichmann et al., encompasses 12 tissues from 38 datasets, forming a meticulously curated cross-tissue database with approximately 3.7 million cells[71]. We extracted 318,426 cells from normal lung samples for our analysis. To differentiate it from another dataset curated by the same group, we refer to this dataset as Teichmann et al.’s 1.
10. Normal samples from human lung immune cells: This dataset profiled human embryonic and fetal lung immune cells using scRNA-seq, smFISH, and immunohistochemistry[72]. For our analysis, we retained normal samples, which include 670,749 cells. As this dataset is also organized by Teichmann et al., we refer to it as Teichmann et al.’s 2.
11. Primary breast tumor atlas: This dataset integrated single-cell RNA sequencing atlas of the primary breast tumor microenvironment containing 236,363 cells from 119 biopsy samples across eight datasets[42]. We utilized it as a case study to explore the outcomes of reference-based CCC analysis using the human breast CCC reference.

#### Simulation and benchmarking design

We designed simulations and benchmarking for method comparison on each dataset describe in the previous section. We considered three distinct set of comparisons:

1. Method validation. We first validated the analytic solution provided by FastCCC with the permutation-based approach CellPhoneDB on the same *CS* used in CellPhoneDB. We used the same log-transformed transcriptomic expression data, the same version of the LRIs database, and the same set of valid LRIs described in the earlier section, for both methods. We conducted the validation experiments using a standard Linux server running Ubuntu 20.04, equipped with an Intel(R) Xeon(R) CPU E5-2620 v2 and 250GB of memory. In the comparison, we kept the parameters used in FastCCC consistent with those used in the default setting of CPDB. For example, FastCCC also used the same mean function for ligand and receptor expression summary statistics, the same minimum function for multi-subunit complexes, and the same arithmetic mean, as used in CPDB. All other CPDB parameters were set to their defaults and we varied the number of permutations used in CPDB (default: 1,000) to examine their influence on CPDB’s final results. In this benchmarking analysis, CPDB results obtained using one million permutations were treated as the ground truth to validate the theoretical derivation and software implementation of FastCCC, as well as CPDB results generated with varying numbers of permutations. We calculated precision and IoU for the results generated by FastCCC and CPDB and assessed the correlation between the p-values produced by the two methods.
2. Simulation comparison. We compared the performance of all methods using the test datasets described earlier. For each dataset, we utilized the raw gene expression count matrix as input and applied a semi-simulated approach, where the ground truth is known, following [16]. We began by randomly shuffling the cell type labels, ensuring that all LRIs were null and no interaction signals were present. We then randomly selected one cell type pair to exhibit LRIs in a specific direction, with ligands in cell type A communicating to receptors in cell type B. For the selected cell type pair, 30 LRIs were randomly chosen from the candidate LRIs database to serve as significant LRIs, where the expression level of cells from the interacting cell type pair were multiplied by a factor of two. Given the inherent sparsity of the data, we identified cells with zero expression counts for either the ligand or receptor within the interacting cell type pair. From these, 60% were randomly selected, and their zero expression values were replaced. The replacement value was computed by first taking the mean of five randomly sampled non-zero gene expression values and then multiplying it by a random factor drawn from a uniform distribution (0.6, 1.2). This adjustment ensured that the ligand and receptor expression levels for the interacting cell type pair were higher than those for a randomly chosen cell type pair. Following these modifications, log1p-normalization was performed, and the normalized counts were used as input for all compared CCI tools (this normalization step was skipped for methods requiring a count matrix as input). Ten simulation replicates were conducted for each dataset. All methods were executed on a high-performance computing cluster with Ubuntu 20.04 and Intel(R) Xeon(R) CPU E5-2683 v3. Tasks were submitted via Slurm, with each node allocating 500 GB of memory. Because some methods rely on built-in LRI databases that are difficult to replace, we applied all methods to the intersection of the LRI database used by each method.
3. Reference-based CCC benchmarking. Here, we evaluated the performance of FastCCC in reference-based CCC analysis using the human CCC reference, comparing it to conventional CCC analysis that solely relies on the user-provided query dataset. Here, we treated the results in the conventional CCC analysis as the ground truth in testing scenarios with biased data subsets.

#### Evaluation metrics

We evaluated the performance of different methods using multiple evaluation metrics. Specifically, based on the ground truth, we first calculated true positive (TP), true negative (TN), false positive (FP) and false negative (FN). Here, TP represents the number of true ligand-receptor interactions correctly detected; FN represents the number of true ligand-receptor interactions not detected; FP represents the number of false interactions detected; and TN represents the number of false interactions not detected. Afterwards, we calculated nine distinct evaluation metrics including accuracy, precision, recall, specificity, macroF1, balanced accuracy, Jaccard index (or intersection over union), Mathew’s correlation coefficient (MCC), and false positive rate (FPR), detailed below:

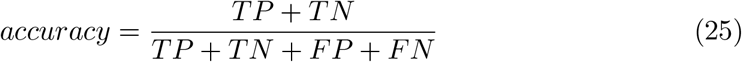

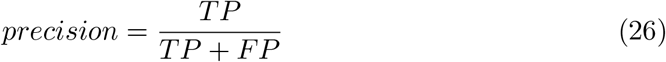

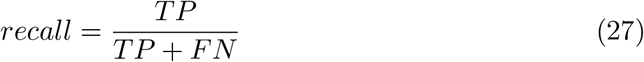

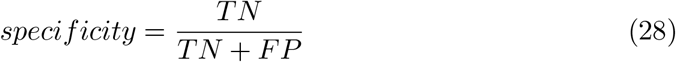

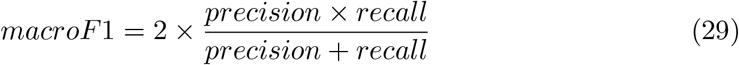

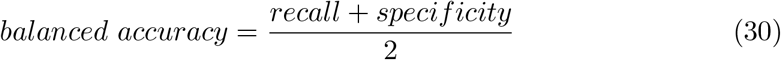

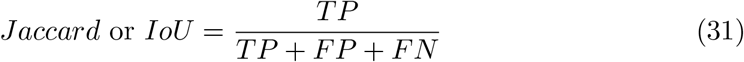

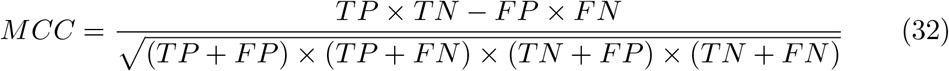

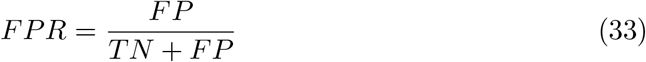

## 5 Data availability

Most of the datasets used for case analysis and comparative experiments in this study were downloaded from the CellxGene[37] platform (https://cellxgene.cziscience.com/). The specific datasets and their sources are as follows:

CPDBTD is available at https://github.com/ventolab/CellphoneDB/blob/v5.0.1/notebooks/data_tutorial.zip.

Ileum is available from https://cellxgene.cziscience.com/collections/ff668d5d-5b3f-49ee-a007-ff0664bf35ec.

Fibroblast dataset is available from https://cellxgene.cziscience.com/collections/a18474f4-ff1e-4864-af69-270b956cee5b.

ECGE is available from https://cellxgene.cziscience.com/collections/cae8bad0-39e9-4771-85a7-822b0e06de9f.

BreastB is available from https://cellxgene.cziscience.com/collections/4195ab4c-20bd-4cd3-8b3d-65601277e731.

Kidney dataset is available at https://explore.data.humancellatlas.org/projects/29ed827b-c539-4f4c-bb6b-ce8f9173dfb7

Angular is available from https://cellxgene.cziscience.com/collections/d17249d2-0e6e-4500-abb8-e6c93fa1ac6f.

Retina is available from https://cellxgene.cziscience.com/collections/5900dda8-2dc3-4770-b604-084eac1c2c82.

GBmap is available from https://cellxgene.cziscience.com/collections/999f2a15-3d7e-440b-96ae-2c806799c08c.

AIDA is available at https://cellxgene.cziscience.com/collections/ced320a1-29f3-47c1-a735-513c7084d508.

HLCA is available at https://cellxgene.cziscience.com/collections/6f6d381a-7701-4781-935c-db10d30de293.

Disease and normal samples of MDCs atlas can be downloaded through https://cellxgene.cziscience.com/collections/3f7c572c-cd73-4b51-a313-207c7f20f188.

LungMAP is available at https://cellxgene.cziscience.com/collections/625f6bf4-2f33-4942-962e-35243d284837

Normal lung samples from HCA is available at https://explore.data.humancellatlas.org/projects/58028aa8-0ed2-49ca-b60f-15e2ed5989d5

Tabula Sapiens is available at https://tabula-sapiens.sf.czbiohub.org/, and we use the normal lung sample.

Two normal lung datasetes organized by Teichmann et al. is available at https://cellxgene.cziscience.com/collections/854c0855-23ad-4362-8b77-6b1639e7a9fc and https://cellxgene.cziscience.com/collections/ec329aed-22bc-4d6e-8935-8282dcb1acac Primary breast tumor atlas is available at https://zenodo.org/records/10672250

For the LRI database, we used version 5.0.0 provided by CellPhoneDB as the primary data source for CCC analysis case studies and reference construction. This database can be downloaded from https://github.com/ventolab/CellphoneDB/blob/master/notebooks/T0_DownloadDB.ipynb. Additionally, version 4.1.0 of the database was also downloaded and used specifically for method comparisons paired with Cell-PhoneDB v4. In the reference-based breast cancer CCC analysis, we also utilized the LRI database from NicheNet v1.1.1 to maintain alignment with the original study. The candidata LRI pairs is extracted from source code at https://github.com/saeyslab/nichenetr/releases/tag/v1.1.1.

## 6 Code availability

FastCCC is freely available as a Python package on github (https://github.com/Svvord/FastCCC) under the MIT license. All scripts used to reproduce all the analysis are also available at the github repostiory.

## Declarations

### Author contribution

X.Z. and S.H designed this project. S.H. and X.Z. contributed key ideas. S.H. conceived the algorithm and developed the software. W.M. and S.H. compared the proposed algorithm with existing methods to evaluate its performance. S.H. performed data analysis and interpreted data. S.H. drafted the manuscript with input from all authors and contributed to the creation and formatting of the figures. X.Z. supervised the project, initiated the work, wrote and edited the manuscript. And all authors read and approved the final manuscript.

## Appendix A Supplementary information

**Table A1.**
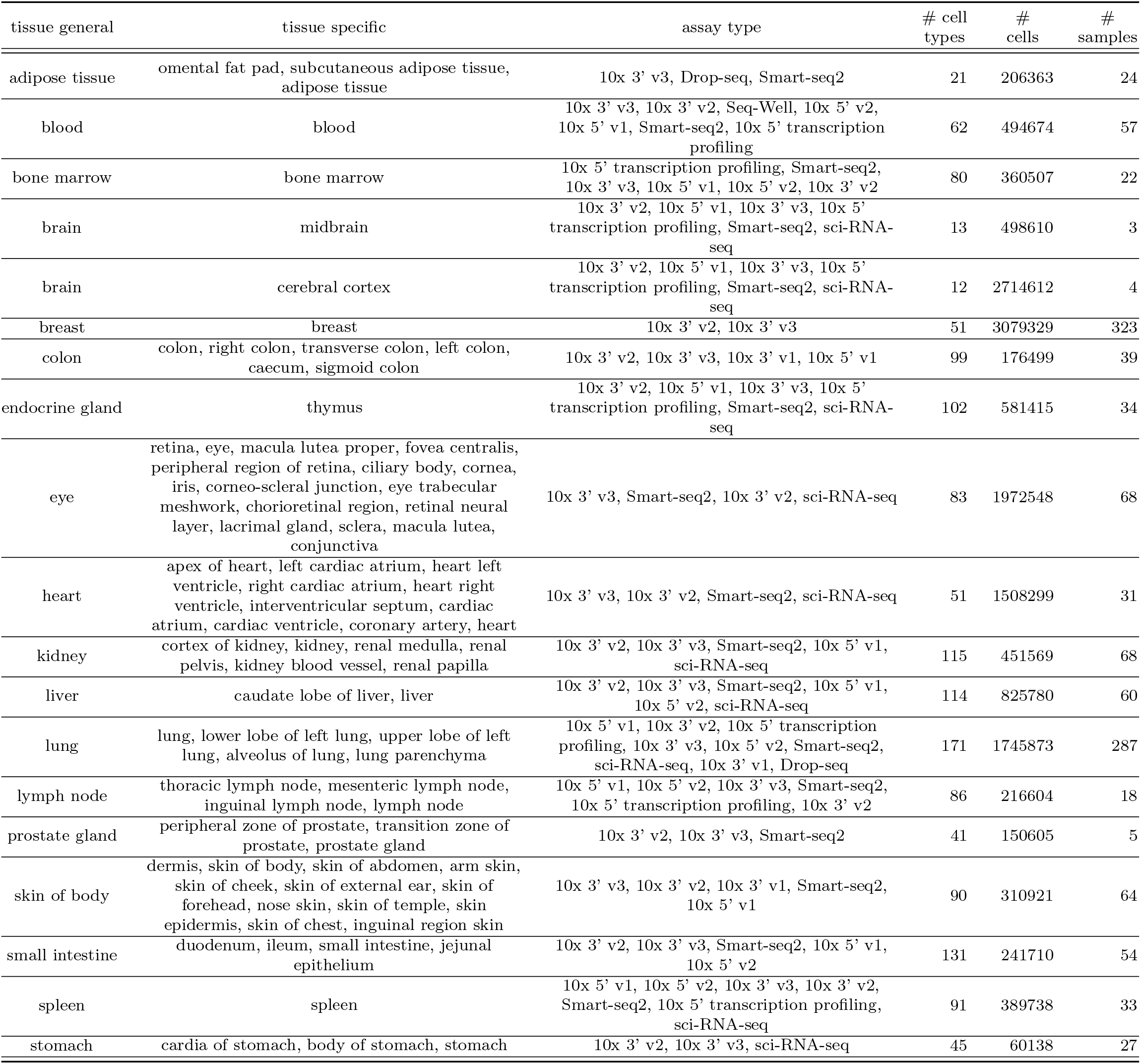
Summary of the Human CCC Reference Panel.

**Table A2.** Summary of the FastCCC strategies.

| cell type | CS | LR sum. | complex agg. | CS dist. | LR dist. | complex dist. |
| --- | --- | --- | --- | --- | --- | --- |
| rare | Eq. 2 | Eq. 4 | Eq. 6 | FFT-based Eq. 8 | FFT-based Eq. 11 | FFT-based Eq. 19 |
| common | Eq. 2 | Eq. 4 | Eq. 6 | FFT-based Eq. 8 | CLT-based Eq. 11 | CLT-based Eq. 19 |
| rare | Eq. 3 | Eq. 4 | Eq. 6 | FFT-based Eq. 10 | FFT-based Eq. 11 | FFT-based Eq. 19 |
| common | Eq. 3 | Eq. 4 | Eq. 6 | FFT-based Eq. 10 | CLT-based Eq. 11 | CLT-based Eq. 19 |
| rare | Eq. 2 | Eq. 5 | Eq. 6 | FFT-based Eq. 8 | Eq. 18 | FFT-based Eq. 19 |
| common | Eq. 2 | Eq. 5 | Eq. 6 | FFT-based Eq. 8 | Eq. 18 | CLT-based Eq. 19 |
| rare | Eq. 3 | Eq. 5 | Eq. 6 | FFT-based Eq. 10 | Eq. 18 | FFT-based Eq. 19 |
| common | Eq. 3 | Eq. 5 | Eq. 6 | FFT-based Eq. 10 | Eq. 18 | CLT-based Eq. 19 |
| rare | Eq. 2 | Eq. 4 | Eq. 7 | FFT-based Eq. 8 | FFT-based Eq. 11 | PMF-based Eq. 21 |
| common | Eq. 2 | Eq. 4 | Eq. 7 | FFT-based Eq. 8 | CLT-based Eq. 11 | CDF-based Eq. 21 |
| rare | Eq. 3 | Eq. 4 | Eq. 7 | FFT-based Eq. 10 | FFT-based Eq. 11 | PMF-based Eq. 21 |
| common | Eq. 3 | Eq. 4 | Eq. 7 | FFT-based Eq. 10 | CLT-based Eq. 11 | CDF-based Eq. 21 |
| rare | Eq. 2 | Eq. 5 | Eq. 7 | FFT-based Eq. 8 | Eq. 18 | PMF-based Eq. 21 |
| common | Eq. 2 | Eq. 5 | Eq. 7 | FFT-based Eq. 8 | Eq. 18 | CDF-based Eq. 21 |
| rare | Eq. 3 | Eq. 5 | Eq. 7 | FFT-based Eq. 10 | Eq. 18 | PMF-based Eq. 21 |
| common | Eq. 3 | Eq. 5 | Eq. 7 | FFT-based Eq. 10 | Eq. 18 | CDF-based Eq. 21 |

Note: “Rare” or “Common” refers to the type of sender or receiver. If the sender and receiver types differ—one being “Rare” and the other “Common”—FastCCC simultaneously applies all strategies used for both cell type categories. “FFT-based” refers to convolution computations performed using Fast Fourier Transform. “CLT-based” denotes approximations derived using the Central Limit Theorem and assuming a normal distribution. “PMF” refers to transforming continuous distributions into discrete probability mass functions for computation, while “CDF-based” involves leveraging the cumulative distribution function of the normal distribution.

**Table A3.** 16 scoring methods implemented in FastCCC.

| Combination ID | Expression Summarization | Multi-subunit Complex | $h(\cdot)$ for $CS$ |
| --- | --- | --- | --- |
| 1 | Mean | Minimum | Arithmetic |
| 2 | Mean | Minimum | Geometric |
| 3 | Mean | Average | Arithmetic |
| 4 | Mean | Average | Geometric |
| 5 | Median | Minimum | Arithmetic |
| 6 | Median | Minimum | Geometric |
| 7 | Median | Average | Arithmetic |
| 8 | Median | Average | Geometric |
| 9 | 3<sup>rd</sup> quartile | Minimum | Arithmetic |
| 10 | 3<sup>rd</sup> quartile | Minimum | Geometric |
| 11 | 3<sup>rd</sup> quartile | Average | Arithmetic |
| 12 | 3<sup>rd</sup> quartile | Average | Geometric |
| 13 | 90<sup>th</sup> percentile | Minimum | Arithmetic |
| 14 | 90<sup>th</sup> percentile | Minimum | Geometric |
| 15 | 90<sup>th</sup> percentile | Average | Arithmetic |
| 16 | 90<sup>th</sup> percentile | Average | Geometric |

**Fig. A1.**
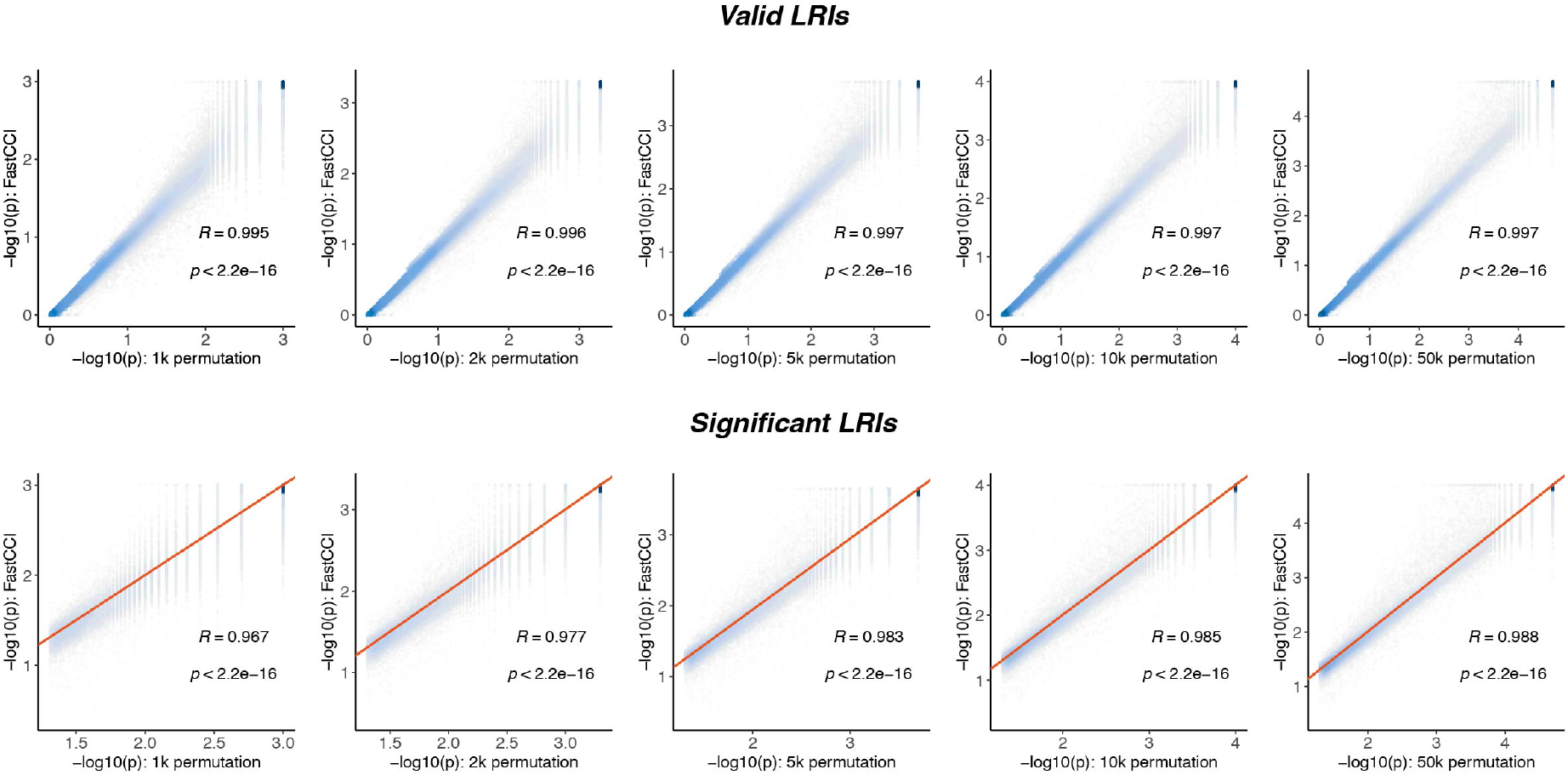
Comparison of LRIs p-values between FastCCC and CPDB across 1,000 to 50,000 permutation tests. Darker colors indicate higher point density. For significant LRIs, the red line represents *y* = *x*. Pearson correlation coefficient and corresponding p-value are annotated.

**Fig. A2.**
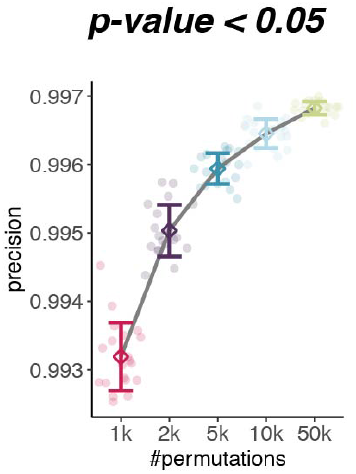
Precision results of significant LRIs (*p*-value ¡ 0.05) between FastCCC and CPDB across varying numbers of permutation tests. Each circle represents an individual experiment (20 repetitions), while squares and error bars indicate the mean and variance.

**Fig. A3.**
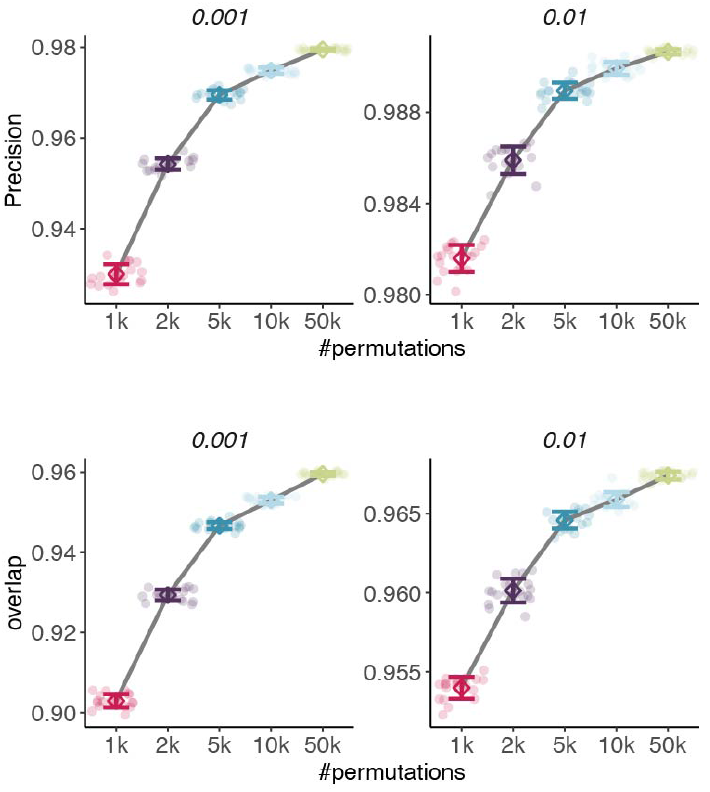
Precision and IoU of significant LRIs identified by FastCCC and CPDB across different permutation tests, evaluated at various *p*-value thresholds.

**Fig. A4.**
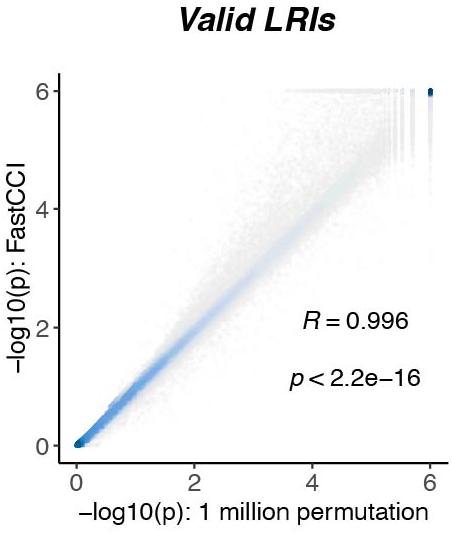
Comparison of LRIs p-values between FastCCC and CPDB under 1 million permutation tests. Each point represents a valid LRI identified by CPDB, with darker colors indicating higher point density. Pearson correlation coefficient and corresponding p-value are annotated.

**Fig. A5.**
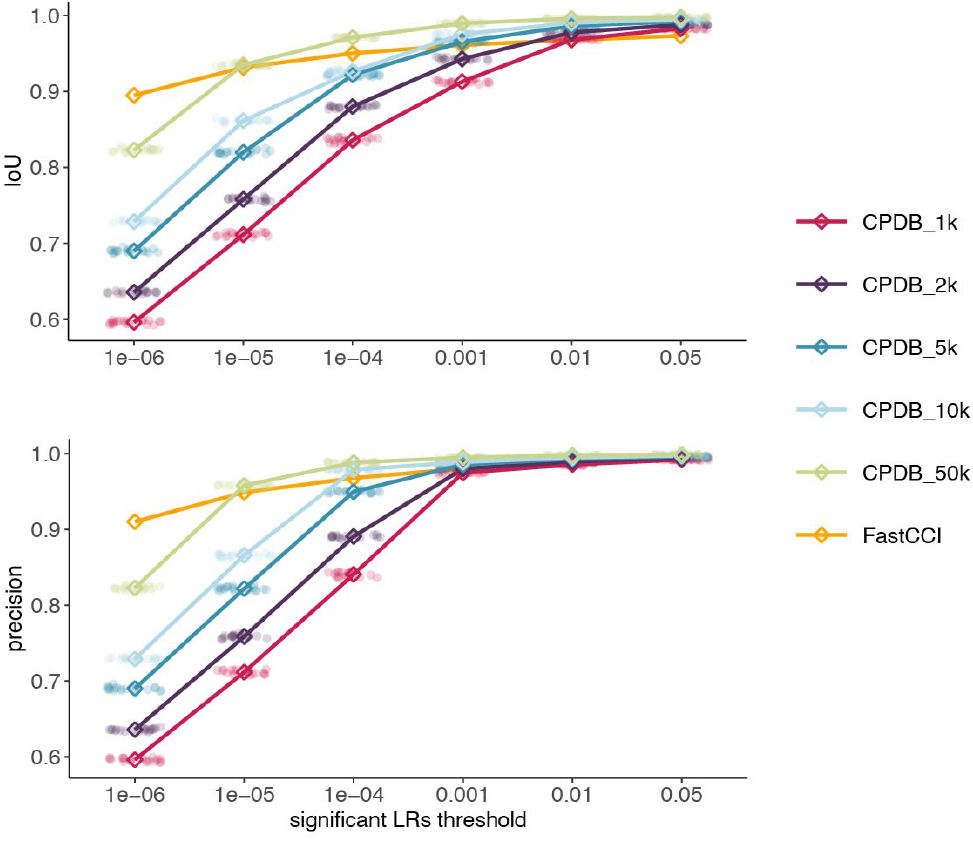
Performance metrics of FastCCC and CPDB at different permutation test numbers compared to the ground truth (1 million permutations) across various thresholds. In general, FastCCC is equivalent to or outperforms CPDB with 10,000 permutations.

**Fig. A6.**
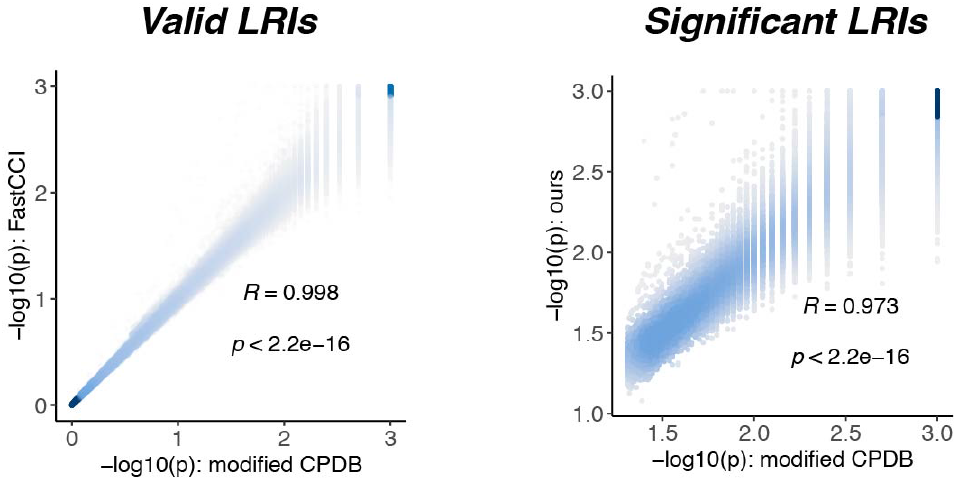
Comparison of modified CPDB and FastCCC results, using the 90th percentile of gene expression as the statistical measure and calculating ligand-receptor communication scores with the geometric mean. Pearson correlation coefficient and corresponding p-value are annotated.

**Fig. A7.**
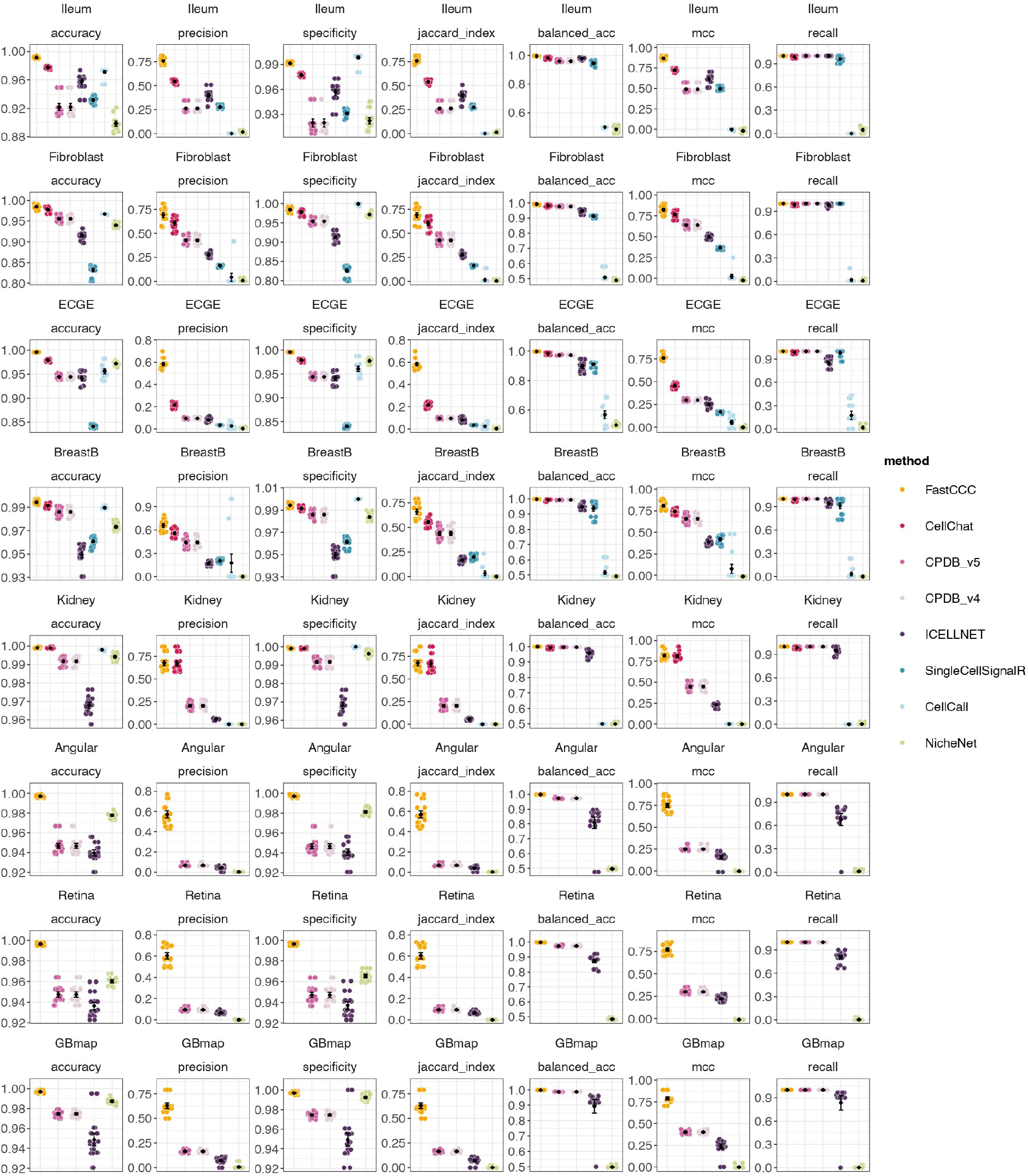
Comparison of FastCCC and other methods across seven metrics on eight test datasets. For all metrics, values closer to 1 indicate better performance. Each point represents an individual experiment, with mean and standard error annotated.

**Fig. A8.**
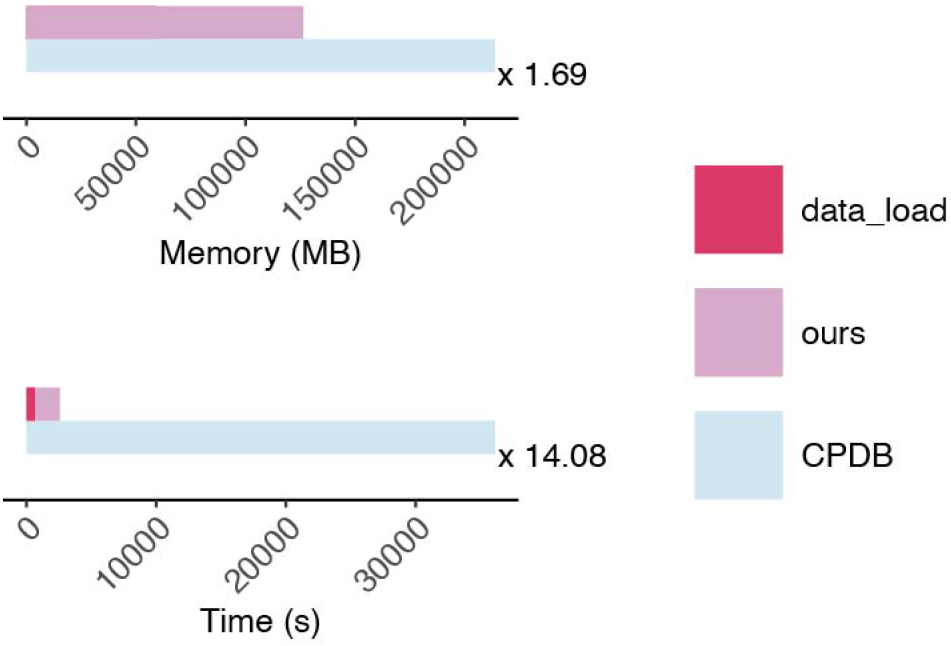
Total runtime and peak memory consumption for running and integrating 16 FastCCC methods on the COVID-19 dataset, compared to CPDB results.

**Fig. A9.**
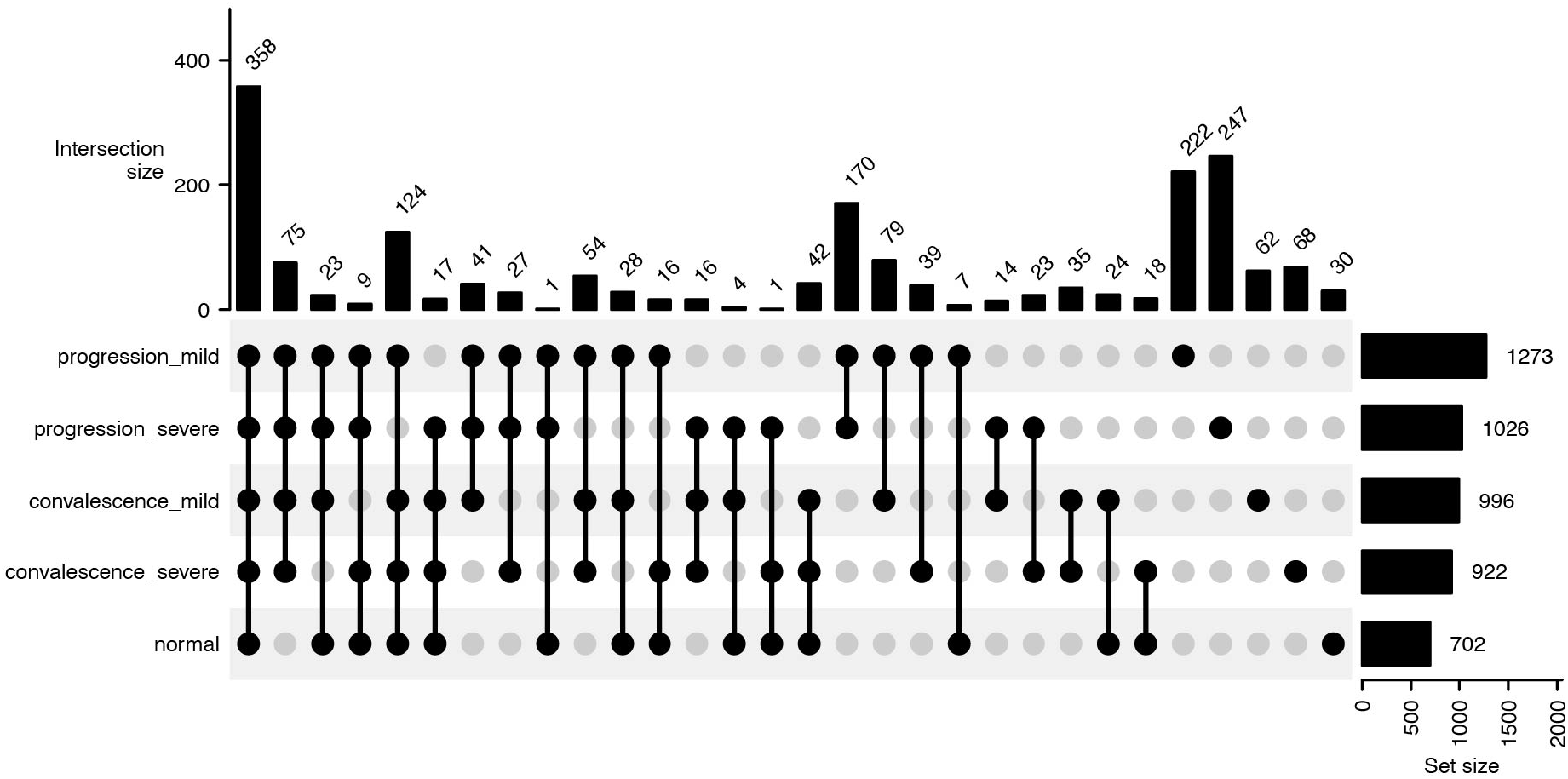
Upset plot showing the intersections of significant LRIs across different groups.

**Fig. A10.**
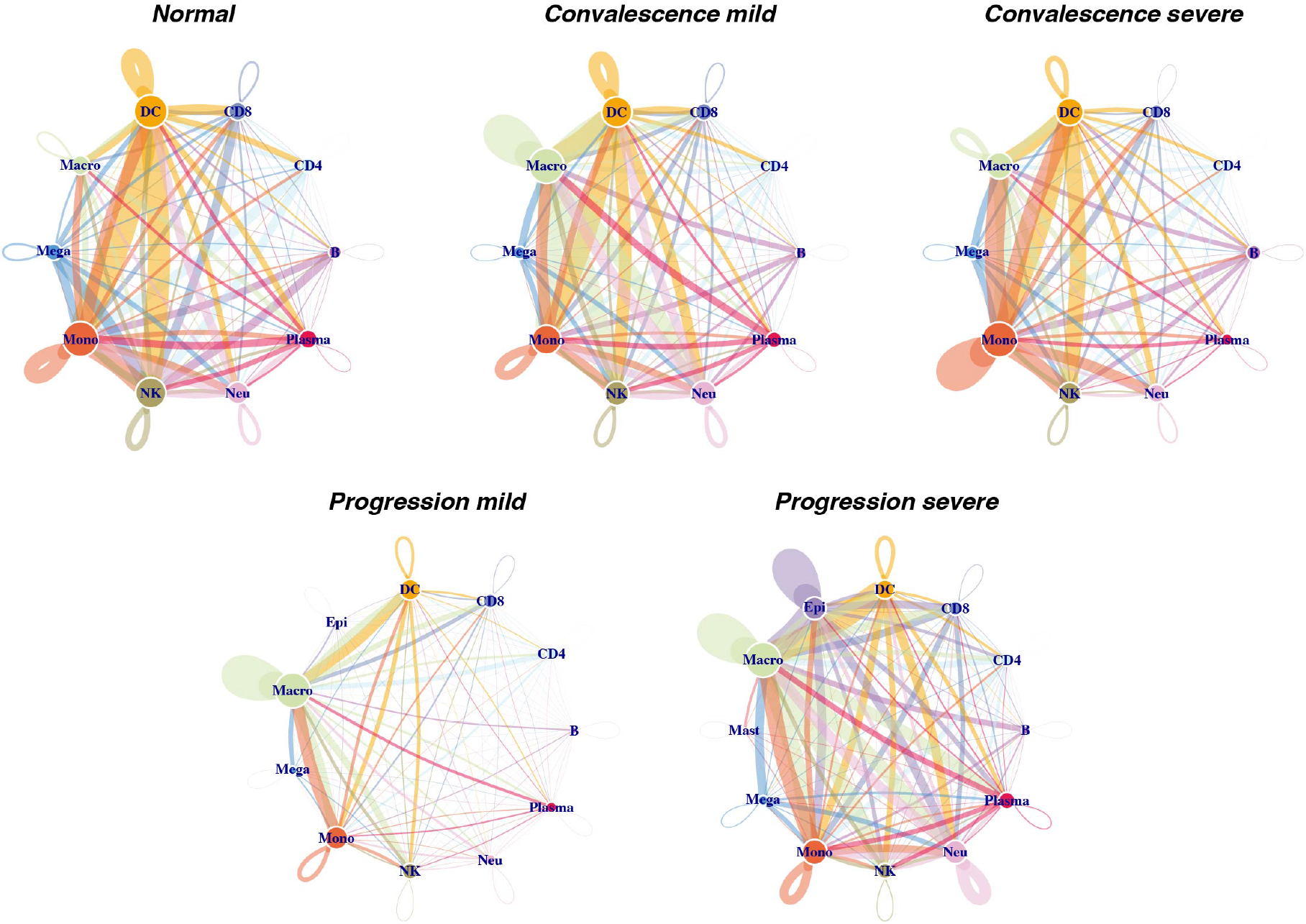
Network plot of distinct groups in the COVID-19 dataset. The size of each circle represents the number of significant LRIs associated with the cell type. The line color indicates the sender cell type, and the line width corresponds to the number of significant LRIs.

**Fig. A11.**
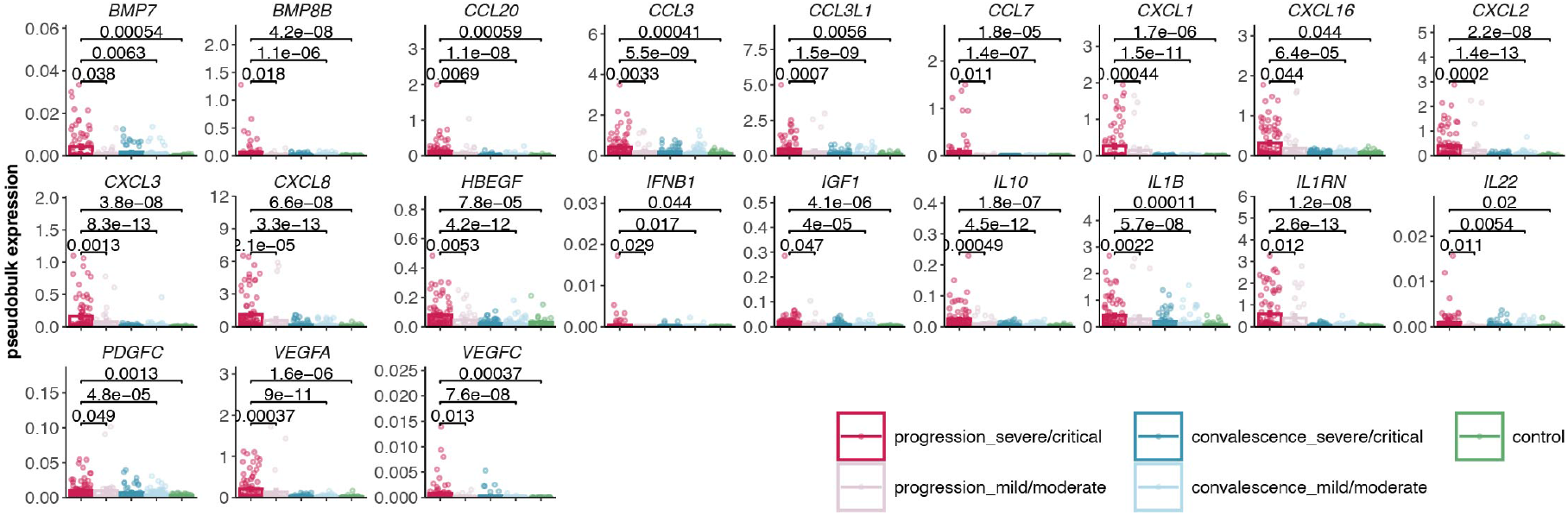
Gene expression of pseudobulk-RNA for differentially expressed cytokine-related genes. Wilcoxon test is performed, and *p*-values are annotated.

**Fig. A12.**
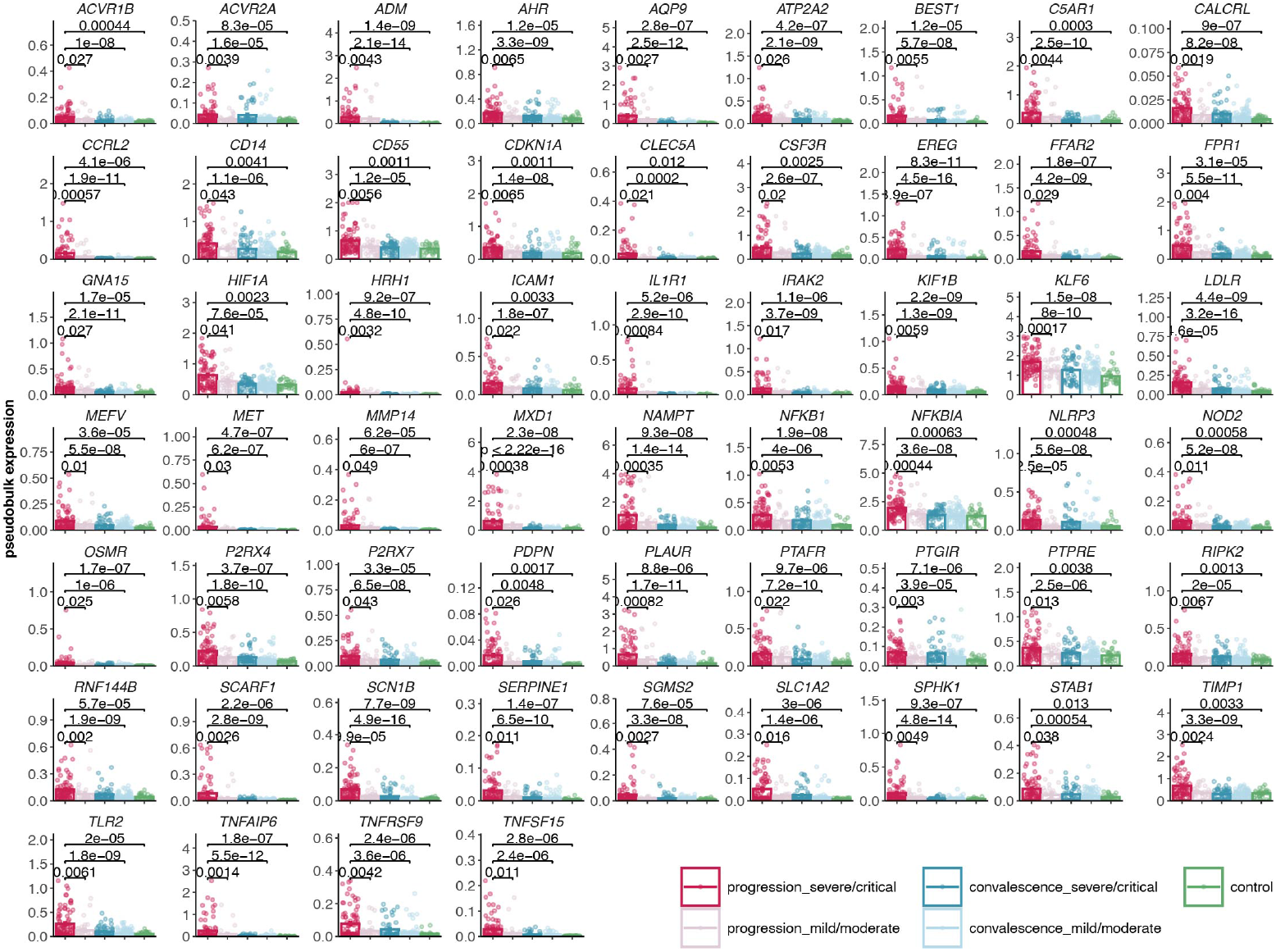
Gene expression of pseudobulk-RNA for differentially expressed inflammation-related genes. Wilcoxon test is performed, and *p*-values are annotated.

**Fig. A13.**
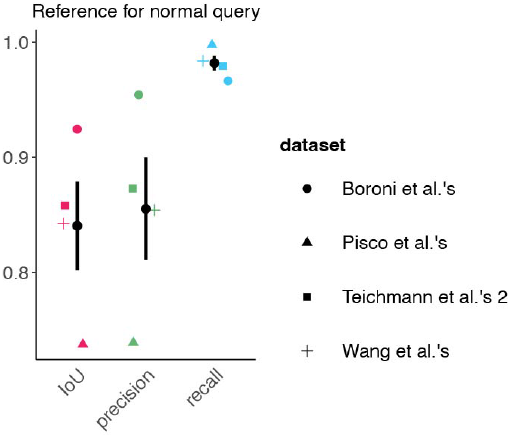
Results of IoU, precision, and recall metrics for normal query datasets using the complete human lung CCC reference panel. Mean and standard deviation are annotated.

**Fig. A14.**
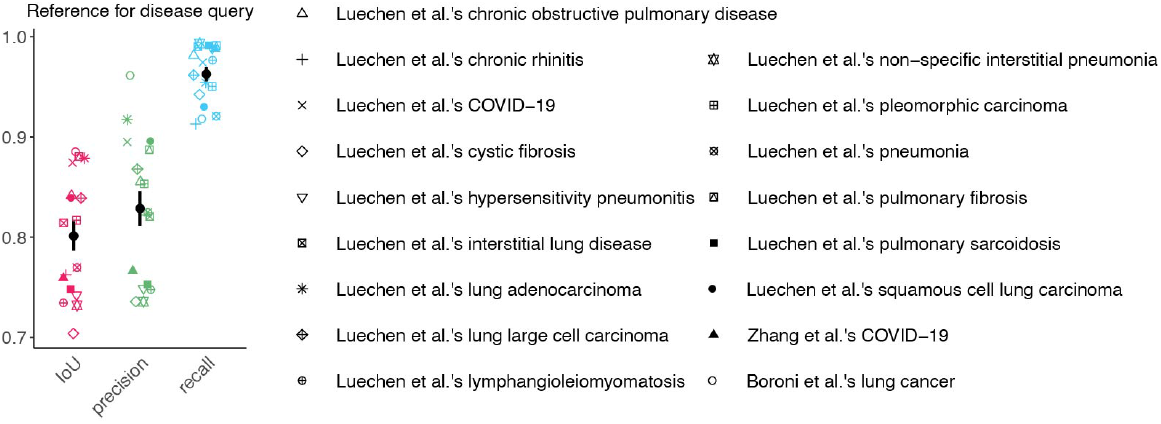
Results of IoU, precision, and recall metrics for disease query datasets using the complete human lung CCC reference panel. Mean and standard deviation are annotated.

**Fig. A15.**
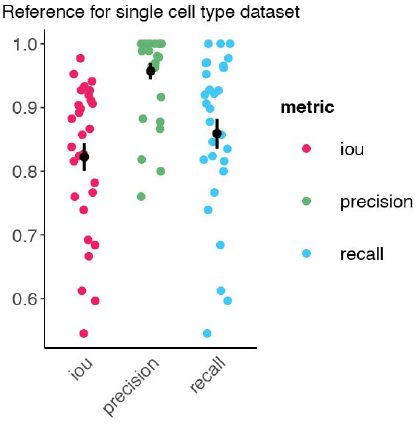
Results of IoU, precision, and recall metrics for the normal lung sub-dataset containing only one cell type. In this scenario, the corresponding FastCCC results from the whole dataset are used as ground truth for those autocrine CCCs. Each point represents a cell type, with the mean and standard deviation annotated.

**Fig. A16.**
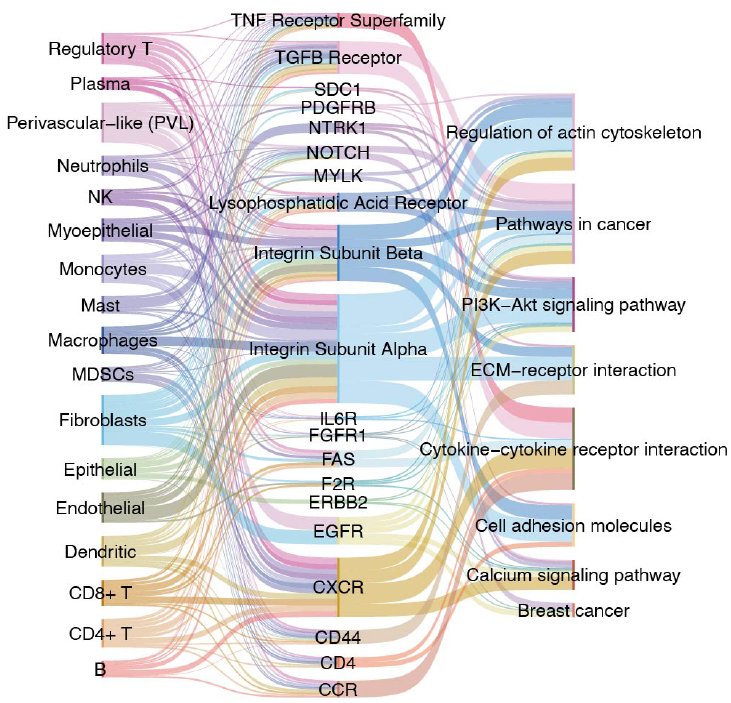
Sankey plot illustrating the relationships between receptors, their receivers, and top enriched KEGG pathways. These receptors frequently appear in significant CCC results and are influenced by a series of upregulated ligands from cancer epithelial cells. They act on various immune cell types, playing regulatory roles in pathways widely reported to be associated with breast cancer.

**Fig. A17.**
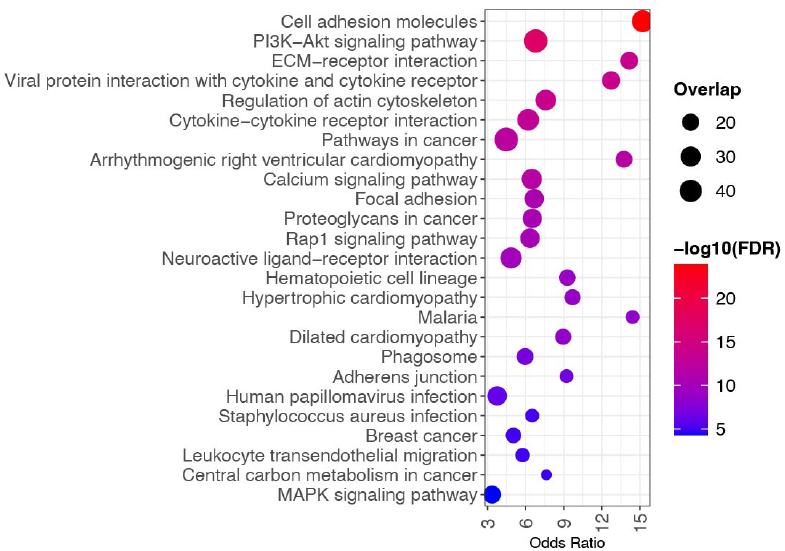
Top 25 enriched KEGG pathways involving receptors influenced by cancer epithelial cells.

**Fig. A18.**
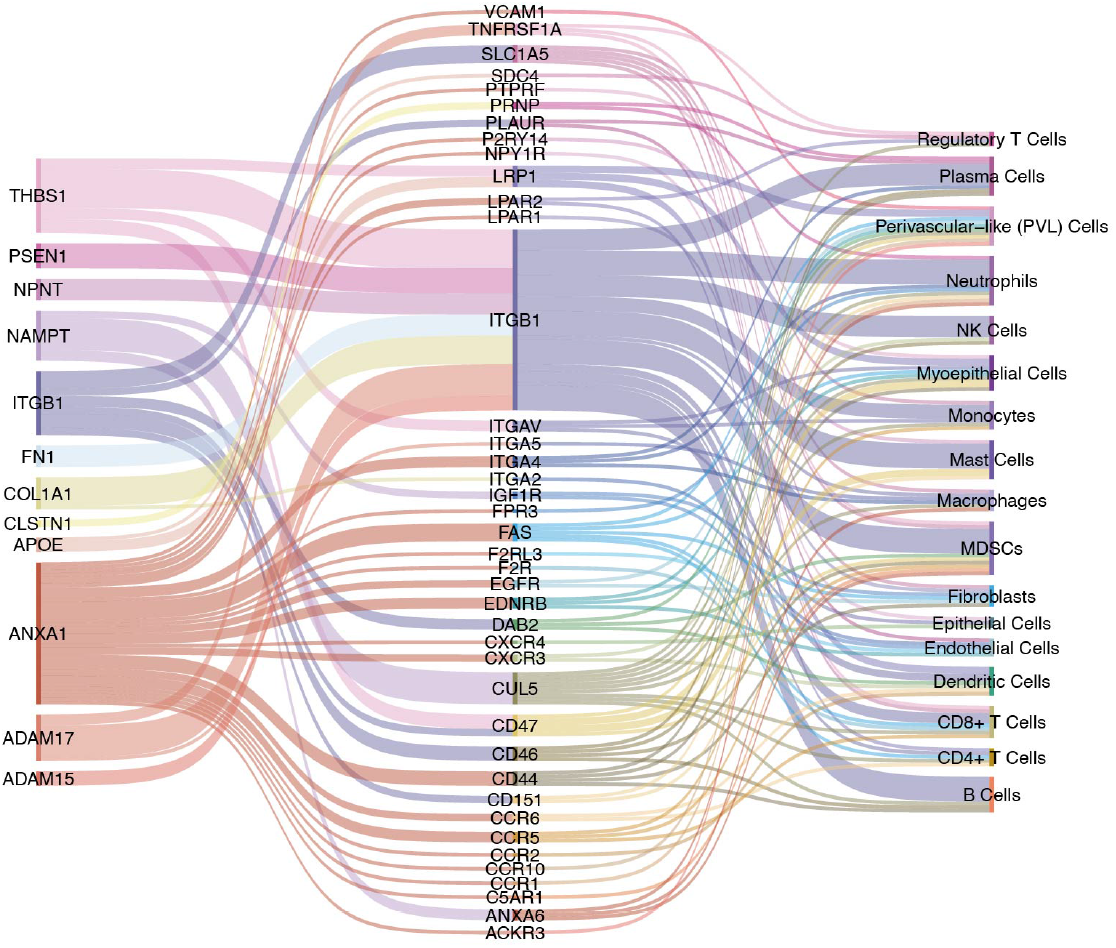
Sankey plot of significantly down-regulated LRIs between cancer epithelial and other cell types. The left column represents ligands in these down-regulated LRIs, the middle column shows the involved receptors, and the right column represents indicates the corresponding receiver cell types. Line reflects the number of involved LRIs, which are widely present in query dataset.

**Fig. A19.**
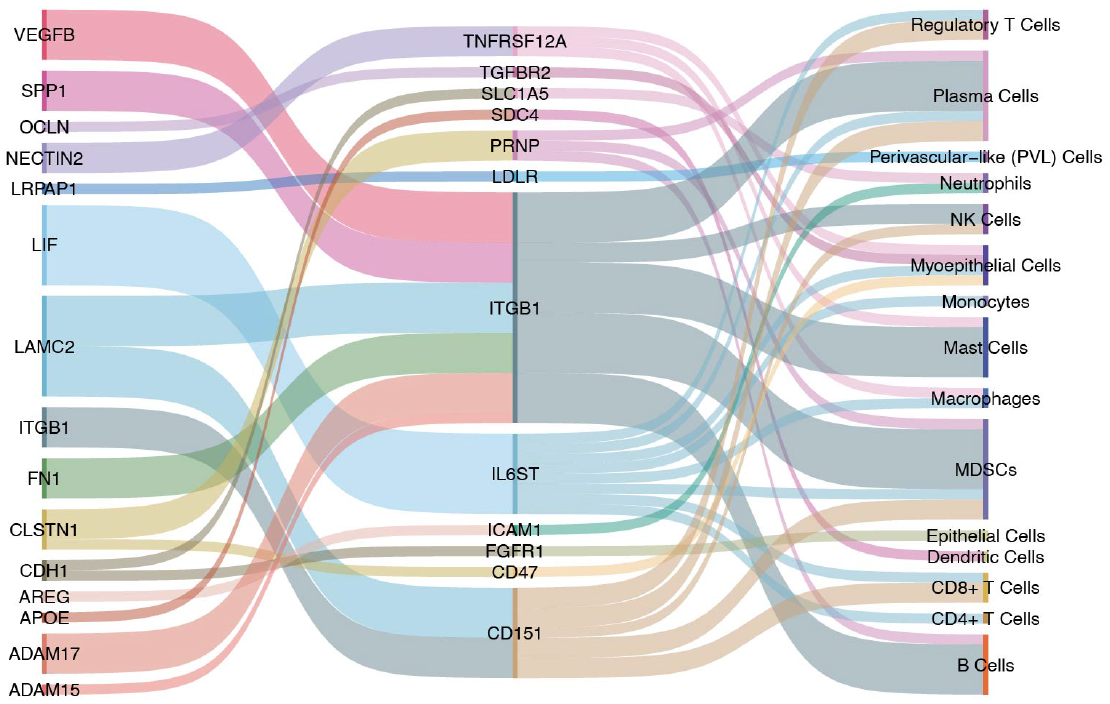
Sankey plot of significantly down-regulated LRIs between cancer epithelial and other cell types in TNBC samples. The left column represents ligands in these down-regulated LRIs, the middle column shows the involved receptors, and the right column represents indicates the corresponding receiver cell types. Line reflects the number of involved LRIs, which are dramatically reduced compared to those in the original dataset.

